# Single cell analysis reveals molecular traits of pediatric lymphoma resistant subclones

**DOI:** 10.64898/2026.04.21.719850

**Authors:** Tracer Yong, Roberta D’Aulerio, Gabriel M. Matos, Marcus R. L. Bezerra, Minghui He, Christian Oertlin, Julien Record, Alexandra Elliot, Anna Kwiecinska, Fredrik Baecklund, Lisa S. Westerberg

**Affiliations:** Department of Microbiology Tumor and Cell Biology, Karolinska Institutet, Stockholm, Sweden; Head and Neck Department, Karolinska University Hospital, Sweden; Department of Oncology-Pathology, Karolinska Institutet Solna, Stockholm, Sweden; Pediatric Oncology, Astrid Lindgren Children’s Hospital, Karolinska University Hospital, Sweden

**Keywords:** pediatric lymphoma, reactive lymph nodes, single cell sequencing, VDJ clonal analysis, spatial transcriptomics, resistant subclone, chemoresistance

## Abstract

The cause of refractory/relapse (R/R) in pediatric lymphoma is unclear. We hypothesized that a stem-like, chemoresistance lymphoma subclone may contribute to R/R. Combining single cell RNA sequencing (scRNA-seq) and immune receptor sequencing (scVDJ-seq) on freshly acquired pediatric non-Hodgkin lymphoma (pNHL n=10), Hodgkin lymphoma (pHL n=5), and 10 reactive lymph nodes from adults or children (a/pReLy), we discovered pediatric-specific progenitor-like lymphocytes, whose cellular program was enriched in pNHL subclones emerging late during clonal evolution, accompanied by loss of immune receptor expression. R-CHOP target gene expression indicated that these subclones may escape first-line treatment, and suggested *MSI2,* an RNA binding protein, as a potential target. In pHL, the progenitor-like program was found in the tumor microenvironment (TME) but not Hodgkin cells which, during relapse, were myeloid-like and accompanied by *CD74*^high^*CCL5*^+^ CD8 T cells. In summary, we discovered R/R associated lymphoma subclones in pediatric lymphoma and potential ways to eradicate them.

**Key messages:**

- Progenitor-like lymphocytes are uniquely found in pediatric reactive lymph nodes, but not in adults.
- Resistant subclones from pNHL acquire progenitor-like program and share *MSI2* expression.
- Relapsed Hodgkin cells are monocyte-like and recruit *CD74*^high^*CCL5*^+^ CD8 T cells.

## Introduction

Pediatric lymphoma accounts for 10-15% of all childhood cancer with ∼60% non-Hodgkin lymphoma (pNHL, B cell ∼40%, T cell ∼20%), and 40% Hodgkin lymphoma (pHL) (*1–3*). Though the difference between pediatric and adult lymphoma is not fully understood, accumulating evidence underscores pediatric lymphoma being a unique entity from its adult counterpart in prevalence, presentation, molecular traits, and treatment response(*4–6*). While treatment strategies differ across lymphoma subtypes, first-line therapy for aggressive pNHL includes agents overlapping with the CHOP regimen (cyclophosphamide, doxorubicin, vincristine, and prednisone) widely used in adults, with the addition of the anti-CD20 antibody rituximab (R) in mature B-cell disease. In Europe, first-line therapy of classical pHL is also CHOP based (*7*), while other regimens are considered outside Europe (https://classic.clinicaltrials.gov/ct2/show/NCT02684708). Although resulting in a cure-rate up to 70-95%, long term adverse effects exist such as infertility, cardiovascular disease, and endocrine conditions (*8*). Moreover, the events of relapses and drug resistance (R/R) are often highly aggressive and associated with poor outcome (*9, 10*). Novel insight into how R/R occurs may benefit future treatment.

Acquisition of stemness is key to cancer progression events such as epithelial mesenchymal transition, metastasis and drug resistance (*11, 12*), and often as a part of cancer evolution (*13*). Although the existence of cancer stem cell is widely acknowledged (*14, 15*), lymphoma stem cells is still debatable (*15*). Most lymphoma, to various degrees, retain their original cellular trait (*16*). For example, clonal expansion of the immune receptor, collectively known as VDJ (Variable, Diversity and Joining) gene segments, and specifically called B cell receptor (BCR) and T cell receptor (TCR), is used for diagnostic in B and T cell NHL lymphoma (*17, 18*). However, to what extent these cell traits are kept at single cell level is unknown. Recent studies identified stem-like single cells in leukemic/lymphoma samples (*19, 20*), supporting the scenario where pre-existing chemoresistant lymphoma subclones being selected by treatment, eventually contribute to R/R (*21, 22*).

Cell type annotation in physiological single cell RNA sequencing (scRNA-seq) data has seen a tremendous improvement over the years (*23, 24*), but precise identification of cancer single cells relies on methods customized to cancer type and conditions (*25*), including unsupervised clustering, marker genes, copy number alteration/variation (CNA/CNV) (*26, 27*), and for lymphoid malignancies specifically, hyperexpanded immune receptor (*28*). Reliable cancer cell annotation with CNV inference allows reconstruction of a cross-sectional phylogenetic tree to reveal key diversification events within a sample (*29, 30*). However, comparisons of evolutionary traits between lymphoma subtypes remain scarce.

In pursuit of a R/R related lymphoma subclone, we here combined scRNA-seq with scVDJ-seq, inferred copy number variation (inferCNV, https://github.com/broadinstitute/inferCNV), single-cell somatic mutations (*31*), and marker gene expression to reconstruct a phylogenetic tree for each lymphoma sample. We discovered a progenitor-like, chemoresistant status as a common trait of pNHL evolution, representing a compartment that had long been over-looked by conventional methods, while Hodgkin cells in R/R became myeloid-like and were associated with CD74^high^CCL5^+^ CD8 T cells infiltration in the TME. These results pave the way to mechanistically understand pediatric lymphoma and ultimately to develop novel treatment that prevent R/R.

## Results

### Characterization of pediatric lymphoid landscape

We prepared scRNA-seq and scVDJ-seq data from fresh lymphoma and reactive lymph node samples collected at Karolinska University Hospital between 2021 to 2025, including non-cancerous pediatric reactive lymph nodes (pReLy, n=8, including 1 revisiting: pReLy-01 and pReLy-01b), adults reactive lymph nodes (aReLy, n=2), Burkitt Lymphoma (pBL; n=3), T cell lymphoblastic lymphoma (pTLBL; n=5), classical Hodgkin Lymphoma (pHL; n=5), anaplastic large cell lymphoma (pALCL; n=1), and primary mediastinal B cell lymphoma (pPMBCL; n=1) (**Figure 1A**, **Table 1**). After quality control (see methods), 21,851 and 60,032 single cells remained for pReLy and aReLy, with mean unique gene count per sample ranging between 2,615∼6,587 and 3,819∼8,297, and mean unique feature count 1,114∼1,987, and 1,375∼2,636, respectively. 21 distinct cell types were found by combining a machine learning algorithm trained by public data sets (*16, 32*) with expert knowledge. To avoid patient specific result in unsupervised clustering, the Ig heavy and light chain loci and TCR alpha/beta loci was excluded from scRNA-seq (*33*).

**Figure 1.**
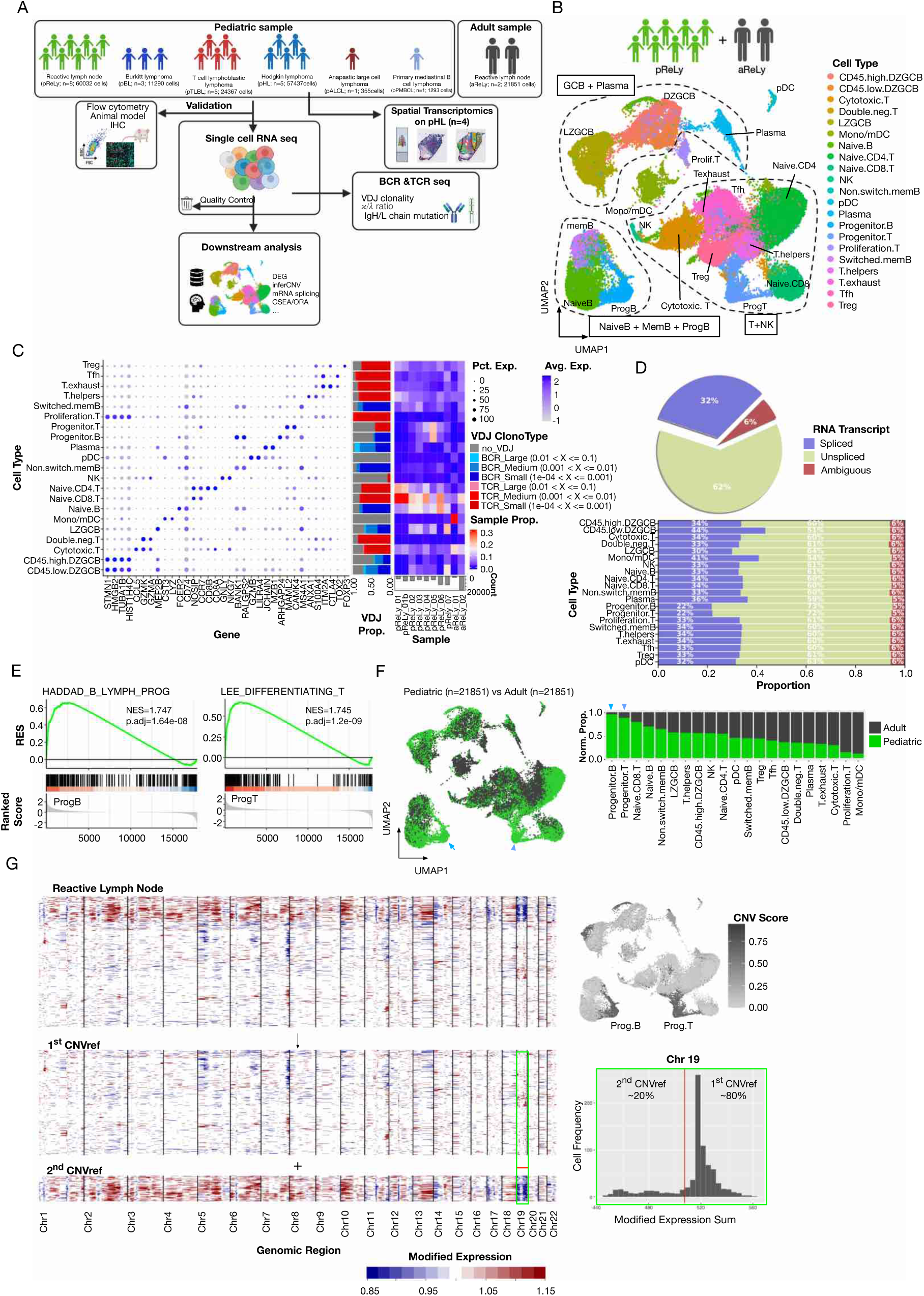
Determining the lymphoid landscape in pediatric specimens: **A.** Schematics of the study. 23 freshly isolated reactive and cancerous pediatric (<18 y.o.) lymph nodes, and two reactive adult lymph nodes were included. Samples were prepared for single cell RNA sequencing (scRNAseq), single cell VDJ sequencing (scVDJseq, BCR and TCR), flow cytometry and, for Hodgkin lymphoma (pHL), spatial transcriptomics. (Schematics created in https://BioRender.com) **B.** Cell type annotation of integrated reactive lymph node (pediatric and adult) represented with 2D UMAP. **C.** (Left) Dot plot highlighting the top 2 marker genes for each identified cell type. Size of the dot indicates percentage of expression, variation in color specifies expression. (Middle) Proportion of VDJ clonotype by cell type. (Right) Proportion of each cell type distribute across all sample integrated in B. **D.** Transcript splicing status by cell types. **E.** Gene set enrichment analysis of progenitor B and T cells against progenitor B and differentiating T cell gene sets, respectively. **F.** (Left) UMAP distribution of adult lymph node cells (n= 21,851 cells) and randomly selected 21,851 pediatric cells. (Right) Proportion of sample source by each cell type. Arrowheads indicate progenitor clusters. **G.** (left) Representative inferCNV heatmap of reactive lymph node sample. Each row represents a randomly selected cell, and each column represents a gene ordered by chromosomes. Color code indicates inferred chromosome gain/loss. (Top right) CNV score distribution across reactive lymphoid landscape. (Lower right) separation of CNV reference based on chromosome 19 expression resulted in two compartments: 20% with high CNV score (2^nd^ CNVref) and the rest 80% (1^st^ CNVref).

To gain insights into how pediatric immune response differ from adults, we first generated a cell atlas of reactive lymph nodes consisting of pReLy and aReLy. Using the uniform manifold approximation and projection (UMAP), three main clusters were observed; cluster 1) T and NK cell cluster included different T cell populations; naïve CD4/CD8, mature helper T cells, cytotoxic T cells, and NK cells, as well as an unique, progenitor-like T cell cluster (ProgT); cluster 2) naïve and memory B cell which also contained a unique compartment termed progenitor-like B cells (ProgB); and cluster 3) germinal center B cells (light zone; LZGCB and dark zone; DZGCB) and plasma cell cluster captured highly proliferative germinal center B cells and proliferating T cells (**Figure 1B** and **Figure S1A**). Smaller clusters included more rare populations such as monocytes/dendritic cells (DC), and plasmacytoid DCs (**Figure 1B**). Marker genes were in line with each mature cell type whereas ProgB/T shared *MAML2*, a *NOTCH* signaling protein (**Figure 1C**). We also observed substantial drop of VDJ expression among ProgB/T as compared to effector cell types (∼40% **Figure 1C**). As expected, we did not observe hyperexpanded VDJ clones in these samples (**Figure 1C**). Interestingly, ProgB/T were found abundant in 6 out of 7 pediatric patients, but underrepresented among adults (**Figure 1C**), suggesting progenitor-like B/T cells are unique to the pediatric lymphoid landscape. Furthermore, ProgB/T cells contained ∼10% more unspliced mRNA (**Figure 1D**) and significantly enriched in progenitor cell gene sets (**Figure 1E**), strongly suggesting a progenitor status. Detailed investigation of ProgB/T cells showed downregulation of translation related genes and upregulation of *NOTCH* signaling gene sets in both (**Figure S1B**), and *WNT* pathway genes in ProgT cells (**Figure S1C**). Sample-size balanced comparison between pReLy and aReLy revealed that pReLys had a larger population of naïve and ProgB/T cells (arrowheads, **Figure 1F**) while adult samples were dominated by mature, effector cells (**Figure 1F**), indicating ProgB/T compartment was associated with an overall naïve, inexperienced immune landscape.

Chromosomal CNV inferred from transcriptome data has emerged as one of the mainstays for single cancer cell detection, for which a reliable reference is essential. We used inferCNV to explore whether the pReLy dataset is suitable for this purpose. We found two distinctive sets of inferred CNV background with the majority (80%) near baseline, termed 1^st^ CNVref, and a smaller set (∼20% of cells) signified by chromosome 19 loss and short arm chromosome 10 gain (2^nd^ CNVref) and high CNV score overall (**Figure 1G**). Interestingly, the CNV score was concentrated in the ProgB/T clusters (Figure 1G), suggesting that the main contributors of 2^nd^ CNVref were ProgB/T cells. This was confirmed by overlapping of up/down regulated genes of ProgB/T cells with the 2^nd^ CNVref gain/loss region (**Figure S1D**), indicating that the 2^nd^ CNVref was transcriptomically driven and indeed represented as a different transcriptomic status. Moreover, the size of 2^nd^ CNVref compartment was proportional to ProgB/T in each sample (**Figure 1C, S1E-M**), agreeing with the link between ProgB/T and 2^nd^ CNVref. In short, we created a reactive, non-cancerous single cell lymphoid atlas and found progenitor-like cells as a unique trait of pediatric immune landscape that need to be considered separately when using as reference in inferCNV.

### A single-cell atlas of pediatric lymphoma reveals origin-specific traits

To gain an overview of our cohort, we created a pediatric lymphoma atlas by integrating pReLy with all lymphoma samples. We found presence of non-cancerous cell types in all lymphoma samples, while lymphoma exclusive clusters fall outside the three main clusters, suggesting abnormal cell status (**Figure 2A**). Breakdown by cell origin showed single cells from B cell originated lymphoma (pBL, pPMBCL) clustered more frequently with B cell than T cells, with pPMBCL higher in myeloid and regulatory T (Treg) cell infiltration (**Figure 2B**, upper left). For pHL, we observed an even distribution between T and B cell clusters but major expansions in the myeloid cluster and lymphoma-exclusive region that linked plasma cells and progenitor T cells (**Figure 2B**, upper right). For T cell originated lymphoma (pTLBL, pALCL), a lymphoma-exclusive cluster adjacent to progenitor T cells was detected (**Figure 2B**, lower right).

**Figure 2.**
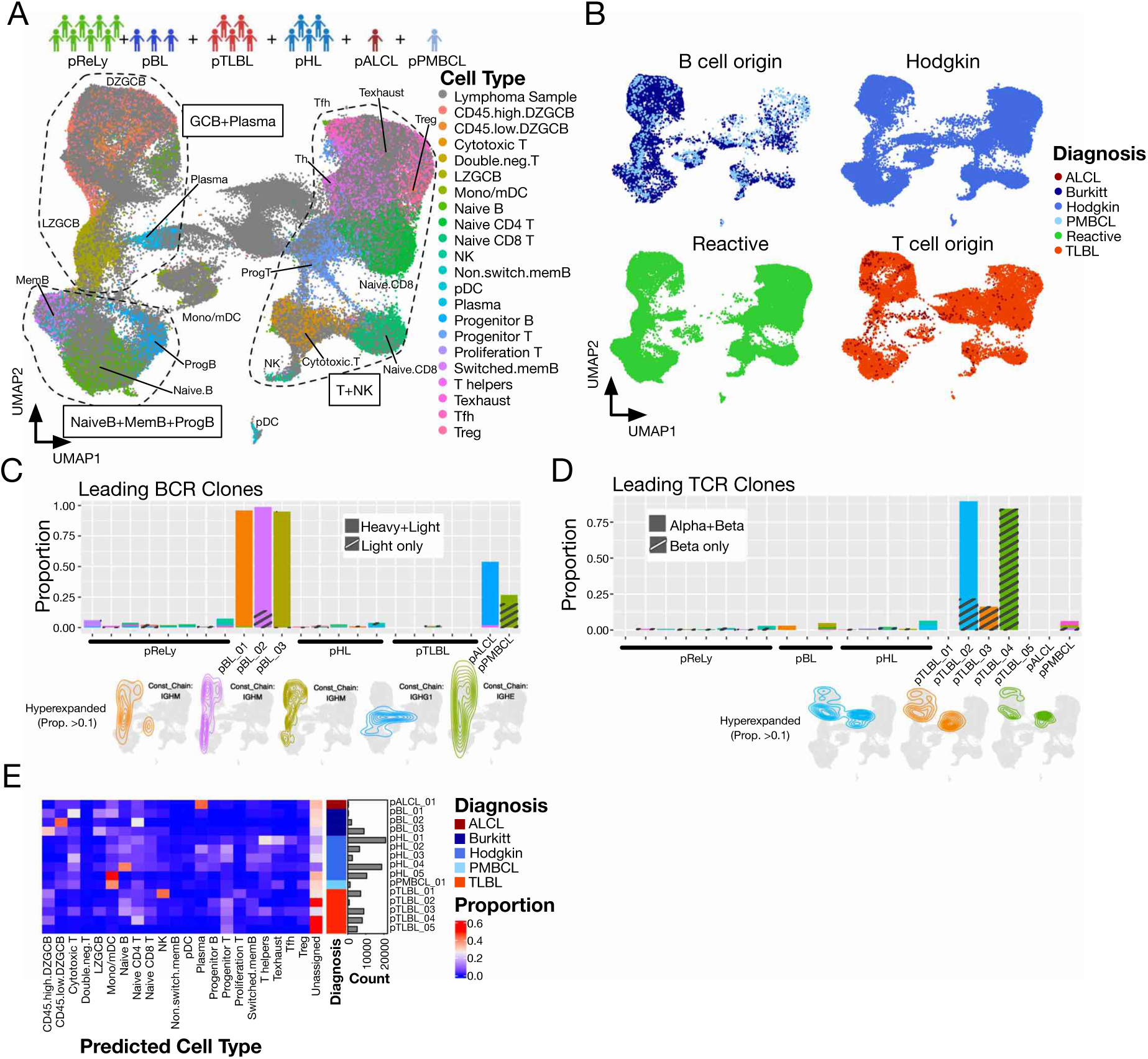
A single-cell atlas uncovers lymphoma origin-specific characteristics. **A.** Integrated scRNAseq UMAP over all pReLy and lymphoma (in grey) samples. **B.** (top) scRNAseq UMAP plotted by pathological origin. **C-D.** Proportion of top 3 BCR and TCR clones per sample. Hyperexpanded BCR clones; pBL_01: IGHV1−46.NA.IGHJ4.IGHM_IGLV2−11.IGLJ2.IGLC2, pBL_02: IGHV3−74.NA.IGHJ6.IGHM_IGLV2−18.IGLJ3.IGLC3, pBL_03: IGHV3−30.NA.IGHJ4.IGHM_IGKV2−24.IGKJ2.IGKC and hyperexpanded TCR clones; pTLL_02: TRAV25.TRAJ45.TRAC_TRBV29−1.NA.TRBJ2−1.TRBC2; pTLL_03: NA_TRBV7−9.NA.TRBJ1−1.TRBC1, pTLL_04: NA_TRBV20−1.TRBD1.TRBJ1−2.TRBC1). (Bottom) Contour plots of hyperexpanded (Proportion > 0.1) immune receptor distribution. Color code matches that of the bar plot. **E.** Proportions of predicted cell types of single cells from lymphoma samples.

To investigate how lymphoma single cells distributed throughout the atlas, we leveraged scVDJ-seq data that identifies lymphoma single cells signified by possessing hyperexpanded immune receptors. For BCR, we found 1 dominant clone (>90%) in each of the 3 pBL samples that transcriptomically span across memory (MemB), light zone (LZGCB) and dark zone germinal center B cells (DZGCB), but not lymphoma-exclusive clusters (**Figure 2C**), suggesting pBL largely retain its canonical B cell program after a single transformation event. A less dominant BCR clone was found in pPMBCL, but with IGHE as constant chain, pointing to its post germinal center origin (**Figure 2C**). Interestingly, a hyperexpanded, class-switched B cell clone was found in pALCL, a T cell lymphoma, indicating effector B cell infiltration (**Figure 2C**).

For TCR, we found hyperexpanded clones existed only in 3 out of 5 of the pTLBL samples, with the other 2 lacking hyperexpanded TCR clones, suggesting that pTLBL transformation occurred during different stages of T cell development in the thymus (**Figure 2D**). Intriguingly, cells from pTLBL occupied a lymphoma exclusive cluster proximal to progenitor T cells and the DZGCB cluster, signifying pTLBL having a unique transcriptome and existence of a proliferating compartment (**Figure 2D**). To investigate transcriptome changes across individual samples, we trained a cell type predictor with pReLy data and applied on all lymphoma samples (**Figure 2E**). High plasma cell content was observed in pALCL, validating effector B cell infiltration observed in Figure 2C. We observed that pBL was predicted to have a high proportion of proliferating cells (DZGCB) followed by pTLBL, whereas pHL had less of DZGCB cells. pTLBL had an increased proportion of naïve CD4 T cells and progenitor T cells and had the largest enrichment of unassigned cells, agreeing with previous observation of a large lymphoma-exclusive cluster. pHL was enriched in a wide range of cell types including naïve B/T, exhausted T, T helper cells, progenitor T cells, and various degree of myeloid cells (**Figure 2E**). In summary, we created a pediatric lymphoma atlas to reveal unique transcriptome and cell type heterogeneity across different origins.

### pBL acquires a progenitor-like, BCR negative program and upregulates MSI2

In search of potential pBL resistant subclones, we integrated 11,290 single cells from pBL samples with pReLy. Most of the single cells from pBL clustered with germinal center (GC) B cells (**Figure 3A**), suggesting Burkitt lymphoma cells largely retain its GC status. This was confirmed by scVDJ-seq data that revealed lymphoma cells, signified by possessing hyperexpanded BCR, spanning across DZGCB, LZGCB, and memB status (**Figure 3B**). Detailed BCR characterization showed deviated light chain usage to either kappa or lambda (**Figure 3C**), and lack of VDJ diversity among dominant clones in pBL samples (**Figure 3D**), with pBL_01 and pBL_03 dominated by a single VDJ and CDR3 sequence (up to 90%) and no diversity found in pBL_02 (VDJ and CDR3 100% identical, **Figure S2**), as confirmed by flow cytometry (**Figure S3A**). This is clearly different from the extensive diversity exhibited by pReLy (**Figure 3C,D**). The evidence indicated that pBL derived from a BCR single cell yet, upon transformation, diversified transcriptomically.

**Figure 3.**
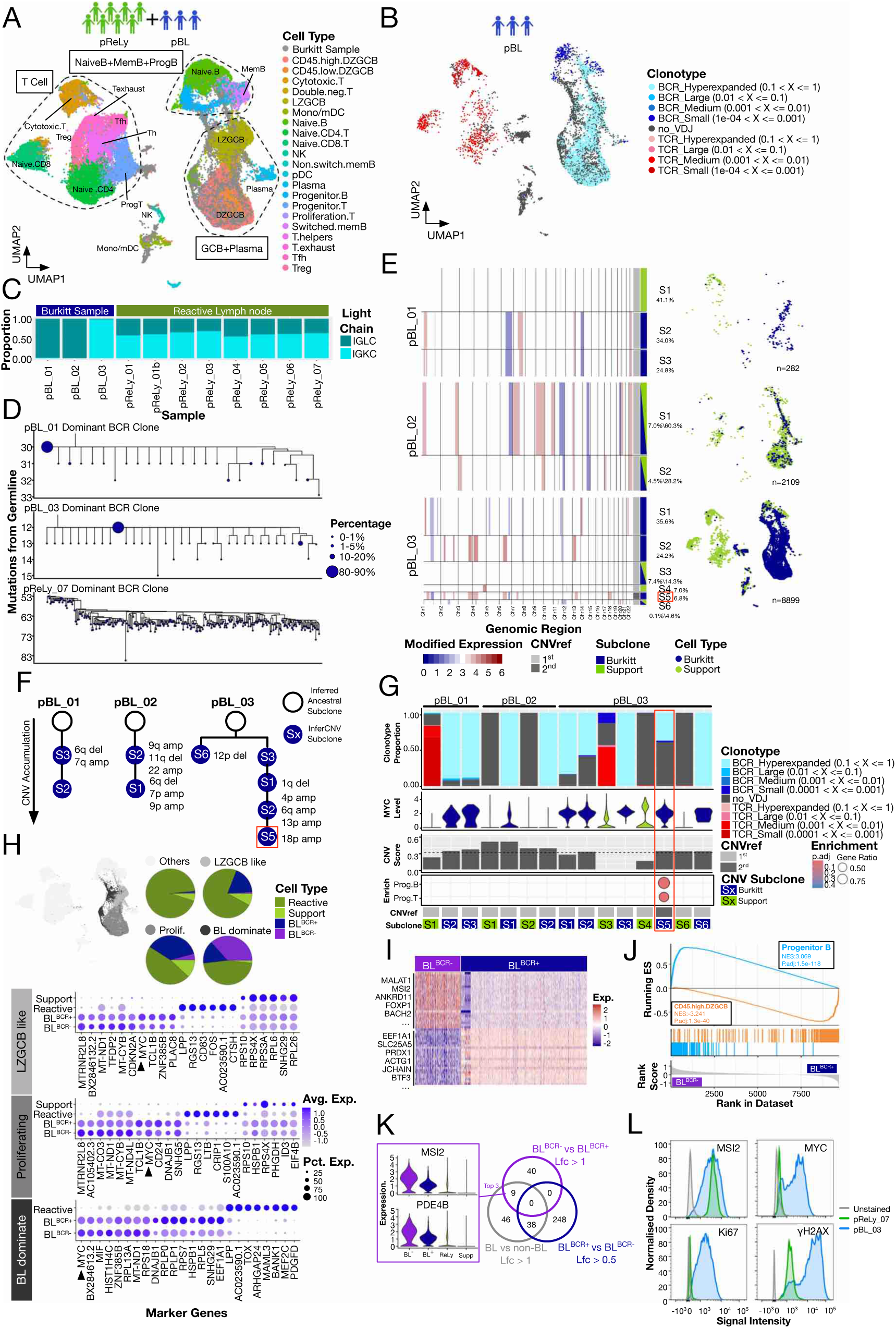
pBL obtains a progenitor-like, BCR negative program and upregulates MSI2: **A.** Integration of pReLy (in color) with three pBL samples (in grey). **B.** BCR/TCR clonotype of pBL single cells. **C.** Proportion of Ig light chain usage (kappa, IGKC or lambda, IGLC) across pBL and pReLy samples. **D.** Phylogenetic analysis of Ig heavy chain CDR3 sequences for hyperexpanded BCR clones from pBL_01 and pBL-03, and a representative leading clone from pReLy_07. **E**. (Left) inferCNV subclustering results of pBLs using the 1^st^ or 2^nd^ CNVref as reference. (Right) UMAP distribution of CNV subclones eventually annotated as “Support” (in light green, defined as non-lymphoma cells from lymphoma samples) or “Burkitt” (in dark blue). **F.** Phylogenetic tree of Burkitt subclones based on CNV accumulation and SNV sharing. **G.** Key characteristics of CNV sublcones from E. (From top) BCR/TCR clonotype proportion, *MYC* expression, CNV score (dotted line indicates 0.31), over representation analysis using progenitor B/T marker gene sets, CNVref used, and eventual Burkitt/Support subclone annotation. **H.** (Top) Three manually curated cell clusters (LZGCB like, Proliferating, and BL dominate) and their composition. Dotplot shows top marker genes of each cell type across these clusters. **I.** Transcriptome difference between Burkitt cells with (BL^BCR+^) or without (BL^BCR-^) functional BCR. **J.** Gene set enrichment analysis of ranked transcriptome comparing BL^BCR-^ to BL^BCR+^ against DZGCB and Progenitor B gene sets. **K.** Venn diagram showing shared upregulated genes across overall Burkitt cells (BL), BL^BCR+^ and BL^BCR-^. Expression of top genes shared by BL and BL^BCR-^ was shown as violin plot on the left. **L.** Representative flow cytometry histograms of *MSI2, MYC, Ki67* and *γH2AX* expression for pReLy_07 and pBL_03.

To capture this diversification after transformation, we subclustered each pBL sample with inferCNV (**Figure 3E-G**). We noticed that CNV for pBL_03 S5 resembled ProgB/T cells in the CNV landscape, meaning the inference could have been skewed by obtaining the progenitor-like program. We revealed the true CNV of this cluster separately with the 2^nd^ CNVref data set (**Figure S1C, S4C**). Combining several key factors describing each subclone including VDJ clonotype, *MYC* expression, CNV score and transcriptome landscape, we were able to annotate these subclones as either Burkitt or Support (defined as non-cancerous cells from the lymphoma sample) and reconstructed phylogenetic trees based on CNV accumulation (**Figure 3F,G**). Since inferCNV is gene count-sensitive only, we validated the phylogenetic tree structures by a sequence-sensitive approach: we detected transcriptome-wide, expressed single nucleotide variation (SNV) at single-cell level via SComatic and calculated Jaccard distances between pBL subclones based on shared SNV. The result indicated a bifurcation during cancer evolution in pBL_03, but not pBL_01 and 02, with S6 isolated from all other subclones (**Figure S4D**), agreeing with our inferCNV-based phylogenetic tree.

Interestingly, we observed a BCR-negative compartment (BL^BCR-^) in pBL_03, a bone marrow sample with the most advanced stage IVB (**Table 1**), increasing in proportion from S1 to S2, eventually acquired progenitor-like program in S5 (**Figure 3G**). This suggests that progenitor-like status was favored under selection pressure. The BL^BCR-^ compartment was negligible in pBL_02, a nodal sample also staged to IVB, suggesting this evolutionary trend could be localized and dependent on environment. Overall, BL^BCR-^ largely retained BL^BCR+^ traits, such as *MYC* expression, across cell clusters capturing most lymphoma cells (black arrowhead, **Figure 3H**). However, when compared to BL^BCR+^, BL^BCR-^ had a distinct transcriptome (**Figure 3I**) and was enriched in Progenitor B cell gene sets (**Figure 3J**), adding to the evidence that pBL_03 was evolving towards a progenitor-like status. Furthermore, comparison between BL^BCR+^, BL^BCR-^, and non-BL cells revealed 9 genes specifically expressed by BL cells with higher expression in BL^BCR-^, among which *MSI2* was the top hit (**Figure 3K**). *MSI2* expression is often linked to hematopoietic and leukemic stem cells (*34–36*). Indeed, *MSI2* protein level exhibited extensive heterogeneity in pBL samples compared to pReLy, confirming existance of a high *MSI2* compartment in pBL (**Figure 3L, S3C**). Additionally, pBL were characterized by high expression of *MYC, Ki67* and *γH2AX* remarking their cancerous nature (**Figure 3L**). Combining these results, we discovered pBL transcriptomically diversify itself after transformation into a progenitor-like BCR-negative subclone in advanced disease.

### Progenitor-like pBL sublones initiate evolution in R/R

We suspected that the BL^BCR-^ compartment represent a stem-like niche that contributes to R/R and applied our pBL pipeline to external pBL scRNA-seq data set (*28, 32*). We detected progenitor-like pBL subclones in 7 out of 12 samples with or without primary refractory (66.7% R/R, 55.6% non-R/R), all harbored considerable proportion of BL^BCR-^ cells (red boxes, **Figure S5A,B**). Upon phylogenetic tree reconstruction, we observed progenitor-like BL subclones among non-R/R samples located terminally at their branches (KC_108, KC_112, KC_113 and KC_115, **Figure S5C**) while those in R/R-related samples were clearly initiating daughter clones (KC_104a and KC_106a, **Figure S5C**), suggesting the maturation of a stem-like niche preceded R/R (**Figure S5C**). Of note, we found KC_101, a non-R/R related sample yet highly expressing the poor prognostic marker *TPM2* (*28*), had a progenitor-like initiating clone (**Figure S5B,C**), indicating this patient was high risk but somehow did not develop R/R. Finally, we observed that *MSI2* had higher expression in the BL^BCR-^ compartment comparing to others, confirming that *MSI2* upregulation is related to R/R in pBL (**Figure S5D**). To sum up, we confirmed *MSI2* upregulation in progenitor-like pBL subclones with external dataset and revealed them often as initiating clones R/R samples.

### Identification of progenitor-like, TCR negative pTLBL compartment

To investigate whether the progenitor-like program also participate in pTLBL lymphomagenesis, we integrated 24,367 pTLBL single cells from 5 samples with the annotated pReLy dataset (**Figure 4A**). CDR3 analysis showed single-dominating TCR clones in pTLBL_02, 03, and 04 but not pTLBL_01 and 05, agreeing with our observation in Figure 2D (**Figure S6**). Despite sharing many normal cell clusters with pReLy, a cell cluster exclusively made up of pTLBL cells was found bridging the T cell and proliferating cluster (**Figure 4A**) and consisted mostly of single cells with hyper-expanded TCR (**Figure 4B**), suggesting this is a lymphoma specific cluster. CNV inference using 1^st^ CNVref revealed progenitor-like subclusters across all pTLBL samples, which we then isolated and processed by 2^nd^ CNVref (**Figure S7A-E**). We annotated inferCNV-derived pTLBL subclusters as either TLBL or Support based on TCR clonality, CNV score, and transcriptome (**Figure 4C-D**). As expected, many lymphoma subclones that required the 2^nd^ CNVref were enriched in progenitor-like program (red boxes, **Figure 4D**), most of which appeared evolutionary terminal (red boxes, **Figure 4E**). Jaccard distance on shared SNV were used to validate major branching events (**Figure S7F**). Of note, S3 in pTLBL_03 was determined to give rise to S4 merely because it had no predicted CNV and, by the assumption of CNV accumulation, is the absolute root of the phylogenetic tree instead of matching S4 CNV pattern (**Figure 4C**). Thus, it is unclear whether pTLBL_03 had an established initiating clone, despite having an advanced disease stage IVA (**Table 1**).

**Figure 4.**
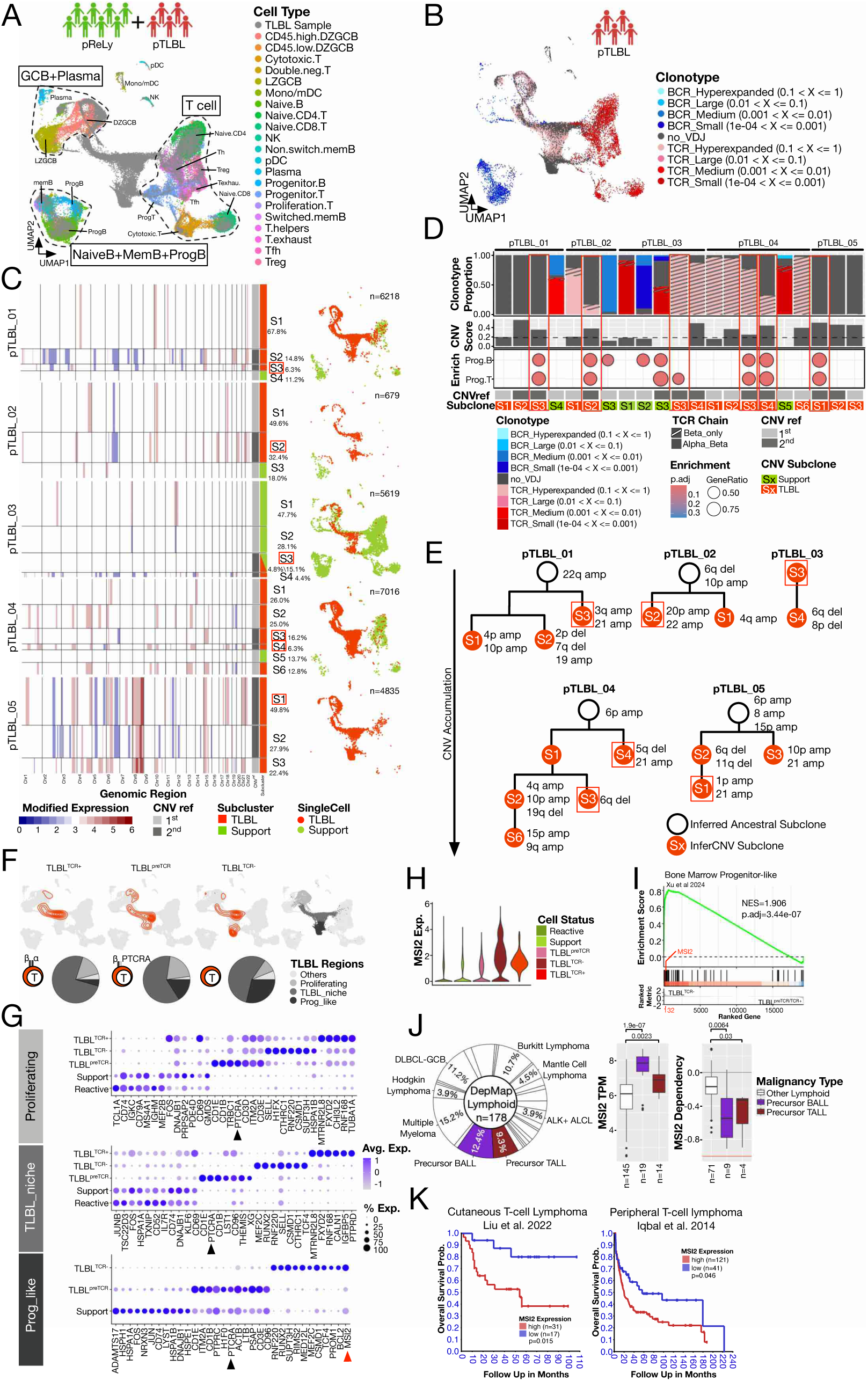
Identification of progenitor-like, TCR negative pTLBL compartment: **A.** Integration of pReLy (in color) with pTLBL samples (in grey). **B.** BCR/TCR clonotype of pTLBL single cells **C**. (Left) inferCNV subclustering results of pTLBLs using the 1^st^ or 2^nd^ CNVref as reference. (Right) UMAP distribution of CNV subclones eventually annotated as “Support” (in light green) or “TLBL” (in orange red). **D.** Key characteristics of CNV sublcones from **C.** (From top) BCR/TCR clonotype proportion, CNV score (dotted line indicates 0.19), over representation analysis using progenitor B/T marker gene sets, CNVref used, and eventual TLBL/Support subclone annotation. **E.** Phylogenetic tree of pTLBL subclones based on CNV accumulation and SNV sharing **F.** Contour plot of TCR status (TCR_pos, Pre_TCR and TCR_neg) distribution of pTLBL single cells and their composition of manually curated cell clusters (Proliferating, TLBL_niche and Prog_like). **G.** Top marker genes of pTLBL (by TCR status), Support and Reactive cells across manually curated cell clusters in **F**. **H.** MSI2 expression across all cell compartments. **I.** Enrichment analysis of bone marrow progenitor-like cells (BMP) related to TLBL^TCR-^ and TLBL^TCR+^ cells. **J.** (left) Cancer type composition of all lymphoid origin cell lines on the Dependency Map (DepMap 2025 Q2). Cell lines annotated as precursor BALL and TALL (based on Expasy Cellosaurus) shown in colors. (right) MSI2 expression and dependency score by precursor status. **K.** Survival curve of closely related T cell lymphoma types by MSI2 expression level.

In pTLBL_03 and pTLBL_04, the hyperexpanded TCR clones consisted of only TCR beta chain (**Figure 4D**). This could not be explained by preferential beta chain coverage over alpha chain during scVDJ-seq as the preference occurs at a limited extent (*37*). Given that the TCR beta chain recombination precedes that for the alpha chain (*38*), it is likely our pTLBL samples originated from different T cell developmental stages. Specifically, pTLBL_01 and 05 were TCR negative upon transformation while pTLBL_02 expressed canonical TCR and pTLBL_03 and 04 expressed preTCR. Thus, we compartmentalized lymphoma cells based on their TCR status: TCR positive (TLBL^TCR+^), Pre-TCR (TLBL^preTCR^) and TCR negative (TLBL^TCR-^). We also manually curated three cellular clusters covering the lymphoma specific region (TLBL Regions, **Figure 4F**). Transcriptomically, all three categories shared the region bridging T helper cells and proliferating cells (TLBL_niche, **Figure 4F**), suggesting that the equilibrium between T cell identity and proliferation is essential to lymphomagenesis. However, in the TLBL^preTCR^, a cluster located towards the progenitor-T cell cluster was observed, which further became a separate cluster in TLBL^TCR-^ (Prog_like, **Figure 4F**), indicating increased representation of progenitor-like program in samples with undifferentiated origin. To validate these findings, we compared top marker genes of lymphoma cells from all compartments across the three manually curated regions (**Figure 4G**). Pre-TCR alpha chain (*PTCRA*) was consistently found among top marker genes for lymphoma cells from TLBL^preTCR^ (black arrowheads, **Figure 4G**), confirming usage of pre-TCR at the moment of transformation. Secondly, lymphoma cells from all three categories had distinct marker genes that pointed to different cellular status. For example, *THEMIS*, a protein critical to T cell development (*39*), was found in TLBL^preTCR^, whereas *RUNX2*, a marker for poor prognosis in T-ALL (*40*), was found in TLBL^TCR-^. For TCR_TLBL^TCR+^, we found *FXYD2*, a gene described in intra-thymic progenitor T cells (*41*) and in CAR-T cells (*42*). Lastly, we identified *MSI2* as one of the top marker genes for TLBL^TCR-^ in the Prog_like cluster (red arrowhead, **Figure 4G**), indicating a TCR negative, progenitor-like *MSI2* high compartment also exist in pTLBL. This is confirmed by comparing *MSI2* expression across all compartments directly (**Figure 4H**).

Given that the TLBL^TCR-^ compartment was also present in progenitor-like TLBL subclones among samples with functional TCR or preTCR at transformation (pTLBL_02 and pTLBL_04, **Figure 4D**), we speculated that TLBL^TCR-^ compartment, like its BL^BCR-^ counterpart in pBL, is responsible of progenitor-like program uptake and is essential to pTLBL progression regardless of TCR status at the moment of transformation. To this end we compared TLBL^TCR-^transcriptome to the other two compartments from the TLBL_niche defined in Figure 4F. We detected enrichment of a bone marrow progenitor (BMP)-like gene set (**Figure 4I**) that has been reported to promote R/R in pediatric T cell acute lymphoblastic leukemia (*43*), a closely related hematologic malignancy, suggesting that the TLBL^TCR-^ compartment is related to R/R. Moreover, *MSI2* was among the BMP-like gene set and ranked 132^nd^ in the full TLBL^TCR-^transcriptome, leading us to hypothesize *MSI2* plays an important role in the progenitor-like BL^BCR-^ and TLBL^TCR-^ lymphoma cells. To gain functional insight of *MSI2*, we categorised all hematologic cancer cell lines on the Dependency Map (*44*) by Expasy Cellosaurus (*45*) into either precursor B cell (n=19), precursor T cell (n=14), or other lymphoid malignancies (n=145) and found malignancies with precursor status had significantly higher expression and dependency on *MSI2* (**Figure 4J**). Lastly, we utilised the survival analyses data base R2 (https://r2.amc.nl/) and detected that *MSI2* expression significantly affected patient survival in two types of T cell lymphoma (*46, 47*) (**Figure 4K**). To summarise, we identified a progenitor-like TLBL^TCR-^ compartment in pTLBL that was associated with R/R and shared high *MSI2* expression with BL^BCR-^.

### Progenitor-like lymphoma cells are potentially chemoresistant

To further explore how the progenitor-like subclones are potentially related to R/R, we integrated lymphoma single cells from either pBL or pTLBL samples (**Figure 5A,C**) and examined expression of (R)-CHOP based therapy related genes, namely, *MS4A1* targeted by rituximab (R) (*48*), *CYP3A5* that activates cyclophosphamide (C) (*49*), *TOP2A* as the main target of doxorubicin (H) (*50*), *TUBB* targeted by vincristine (O) (*51*), *NR3C1* the receptor for prednisone (P) (*52*), and *ABCC1*, also known as multiple-drug resistant protein 1 (MDP1), that mediate efflux of doxorubicin, vincristine and other chemotherapy agents (*50, 53*) (**Figure 5B, D**). Upon integration, BL^BCR-^ cells were transcriptomically distinct from BL^BCR+^ (**Figure 5A**), expressing slightly less R-CHOP targets like *MS4A1* and *TUBB* while expressing higher NR3C1, suggesting the transcriptome change may cause a shift in responsiveness to therapy (**Figure 5B**). Interestingly, BL^BCR-^ cells were found to express more *ABCC1*, a protective factor against doxorubicin and vincristine, showing signs of chemoresistance (**Figure 5B**). Since the TLBL^TCR-^ compartment consists of two clusters in Figure 4F, we separated progenitor-like cells (TLBL_Prog^TCR-^), defined as TCR negative lymphoma cells from Prog_like cluster in Figure 4F, from other TLBL^TCR-^ cells (**Figure 5C**) and found TLBL_Prog^TCR-^ had very low CHOP target gene expression such as *TUBB* and *NR3C1* while expressing, higher *ABCC1*, again suggesting chemoresistance (**Figure 5D**). Of note, *MS4A1* expression was found complementary to that of *TOP2A* and *TUBB* among pBL single cells, covering all areas across BL^BCR+^ (bottom row, **Figure 5B**) while no such pattern was found of in pTLBL (bottom row, **Figure 5D**). This is aligned with current observation that pBL has better prognosis than pTLBL under (R)-CHOP-based regiment (*54, 55*) and prompted us to search for genes expressed complementarily to (R)-CHOP targets. We calculated correlations of gene expression between *TUBB*, the strongest expressed CHOP target, and transcriptome-wide individual genes, revealing top correlated genes such as *TUBA1B*, a close partner of *TUBB*, as well as counter-correlated genes (left panel, **Figure 5E, G**). The top 300 genes negatively correlated to *TUBB* expression were found to enrich different programs across pBL and pTLBL (right panel, **Figure 5E, G**). Specifically, pBL cells with less *TUBB* expression were enriched in a canonical B cell program and small GTPase activity while its counterpart in pTLBL were enriched in cell adhesion and stem cell development, implying different strategies in maintaining chemoresistance. Indeed, expression of the top complementary genes in pBL, *LYN* and *FCRL1*, both linked to BCR signalling (*56, 57*) and in pTLBL, *COL23A1* and *CDH4* that has been associated with anchorage-independent cell proliferation (*58*) and ß-catenin pathway (*59*), respectively, were elevated in the chemoresistant compartments that shared high *MSI2* expression (**Figure 5F, H**), reiterating the importance of *MSI2* in progenitor-like lymphoma compartments.

**Figure 5.**
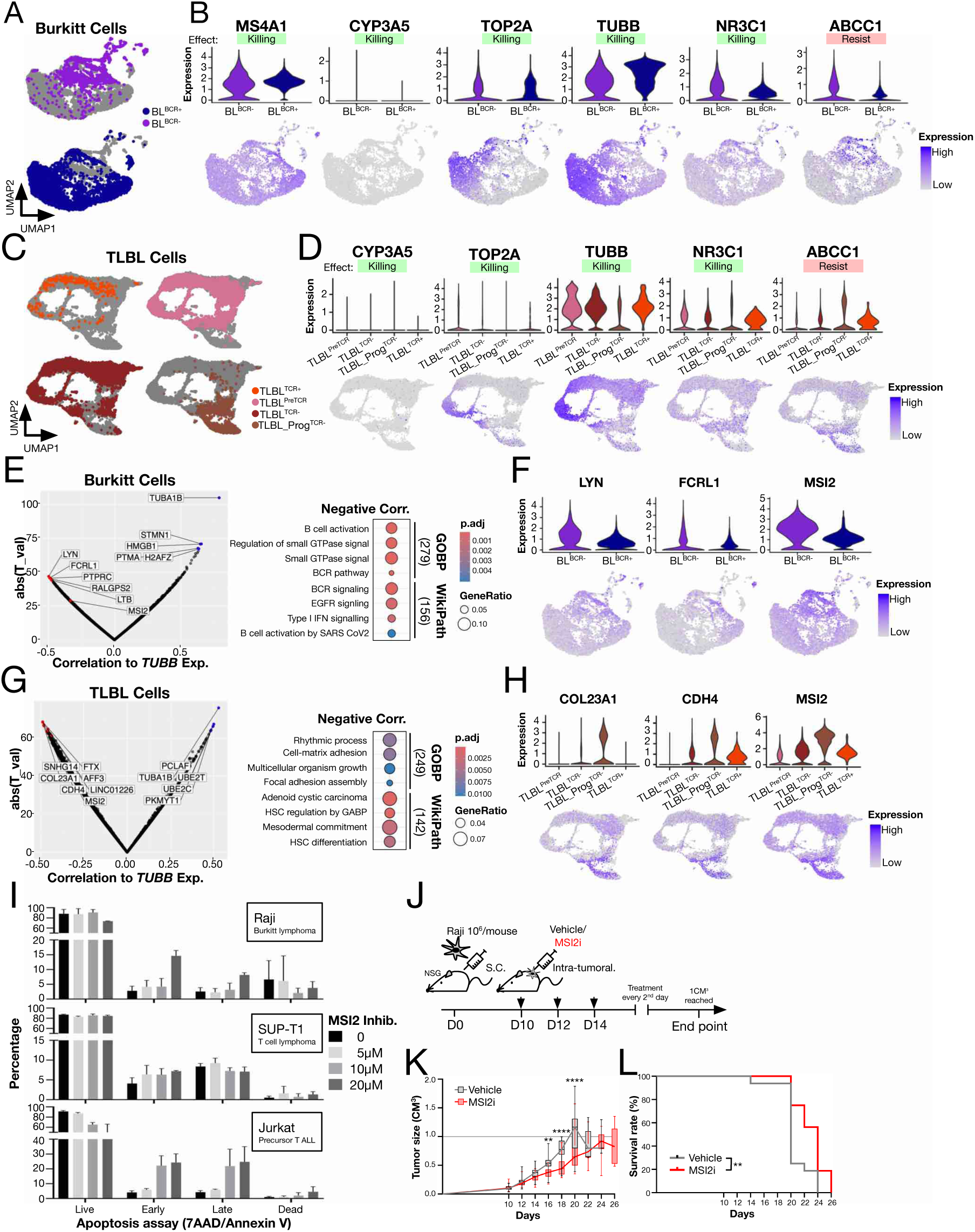
Progenitor-like lymphoma cells are potentially chemoresistance: **A.** Integration of all cancerous pBL single cells. BCR status shown in color. **B.** Expression level of genes involved/targeted by R-CHOP regiment (killing) or related to chemoresistance (resist) in cancerous pBL single cells samples. **C.** Integration of all cancerous pTLBL single cells. TCR status shown in color. **D.** Expression level of genes involved/targeted by CHOP regiment (killing) or related to chemoresistance (resist) in cancerous pTLBL samples. **E.** (Left) expression correlation between TUBB and other genes in pBL. Top positive (in blue) and negative (in red) correlated genes were highlighted together with *MSI2*. (Right) top enriched terms (gene ontology biological process, GOBP; WikiPathway, WikiPath) for top 300 genes negatively correlated with *TUBB* expression. **F.** Highlighted gene expression negatively correlated with *TUBB*. **G.** Same as E. but for pTLBL. **H.** Same as in F. but referred to pTLBL cancerous cells. **I.** Apoptosis assay for Raji, SUP-T1 and Jurkat cell lines upon MSI2 inhibitor treatment. Cells were incubated for 48h at designated concentrations. **J.** Schematics of NSG xenograft model utilizing Raji cell line. MSI2 inhibitor was administered intratumorally every 2 days starting from day 10. Experiment end point is represented by tumour reaching 1cm3. **K.** Tumor volume evaluated at each time point. Statistical analysis performed utilizing 2 way Anova - Šídák’s multiple comparisons test (untill D24). **L.** Percentage of survival with and without MSI2 inhibitor treatment referred to **J.**, curve comparison analyzed with Logrank (Mantel-Cox) test.

To validate that *MSI2* potentially serves as a target for progenitor-like lymphoma cells, we used an apoptosis assay to characterize how *MSI2* inhibitor treatment (*60*) affected 3 hematological malignancy cell lines, namely Raji, a Burkitt lymphoma cell line (RRID:CVCL_0511), SUP-T1, a TLBL cell line (*61*), and Jurkat which belongs to precursor T cell leukemia (RRID:CVCL_0065). After 48-hours, Jurkat cells exposed to varying concentration of *MSI2* inhibitor treatment had the lowest viability among all tested cell lines (**Figure 5I**), highlighting that *MSI2* is important for cells with precursor status. While SUP-T1 cells were not affected by *MSI2* inhibition *in vitro*, Raji cells had increased apoptotic cells upon MSI2 inhibitor treatment (**Figure 5I**). We reasoned that the progenitor-like lymphoma compartment eventually develops *in vivo* and plays important role in disease progression. To test the efficacy in vivo, we injected Raji lymphoma cell line subcutaneously in NSG mice and administered *MSI2* inhibitor intratumorally and monitored tumor growth over time (**Figure 5J**). We observed that Raji lymphoma grew significantly slower after *MSI2* inhibition (**Figure 5K**) and was associated with longer survival time (**Figure 5L**), suggesting benefit of targeting the *MSI2*-high compartment in BL. Collectively, we discovered progenitor-like compartments in pBL and pTLBL were chemoresistant while shared *MSI2* as vulnerability.

### Intratumoral heterogeneity across pHL tumor microenvironment (TME)

To investigate whether progenitor-like program plays a role in pHL, we performed spatial transcriptomics on 4 pHL tissues. Dimension reduction and nearest neighbor clustering identified 15 transcriptomically distinct clusters across all 4 samples. We then applied cell type specific module scores, namely, Hodgkin (*TNFRSF8*, *TXN*, *CCL17*) (*62–64*), histiocyte (*C1QA, LYZ, S100A8*) (*65–67*), M2-like macrophage (*CCL18, MMP9*) (*68, 69*), and B cell (*CD19, MS4A1*) (*70, 71*) to each cluster. This revealed that cluster 3 and 9 were dominated by B cells, whereas several clusters had high histiocyte (macrophage) activity, among which cluster 11 seemed to adopt a M2 macrophage-like program (M2-Mq). Hodgkin cell activity was almost exclusively found in cluster 10, pointing to similar pHL niche composition throughout samples (**Figure 6A**). Regarding heterogeneity of the samples, we observed pHL_02 was voided of 7 out of 15 clusters from the integration and without any cluster unique to it, strongly suggesting lack of intra-tumor heterogeneity and low quality. This is confirmed by quality parameter and expert knowledge (**Figure S8A**). By investigating spatial arrangement of each integrated cluster, we found cluster 3 (B cell) and 11 (M2-Mq) consistently occupied opposite side of cluster 10 (Hodgkin), forming a 3-layered structure that can be observed across all high-quality samples (left panel **Figure 6B, Figure S8B**). This indicated their distinct but universal roles in lymphomagenesis. To validate cluster 10 as the Hodgkin niche, we calculated module score using gene panels curated by pathologist (A.K.) for HRS cell identification and TME characterization, namely, Hodgkin inclusive: *TNFRSF8* (CD30), *FUT4* (CD15), *IRF4*; Hodgkin exclusive: *MS4A1* (CD20), *CD79A, CD19, SLC22A2, POU2AF1*; and T cell exhaustion: *CTLA4, LAG3*. We observed that the Hodgkin inclusive score not only spatially complemented Hodgkin exclusive score, but also co-localized with the T cell exhaustion score, agreeing with current knowledge on exhausted T cell infiltration in the Hodgkin niche (*72*) but avoiding other parts of the TME (right panel **Figure 6B**). Representative genes for M2-Mq (*MMP9*), Hodgkin cells (*TXN*), and B cells (*MS4A1*) validated the spatial arrangement observed in Figure 6B (**Figure 6C**).

**Figure 6.**
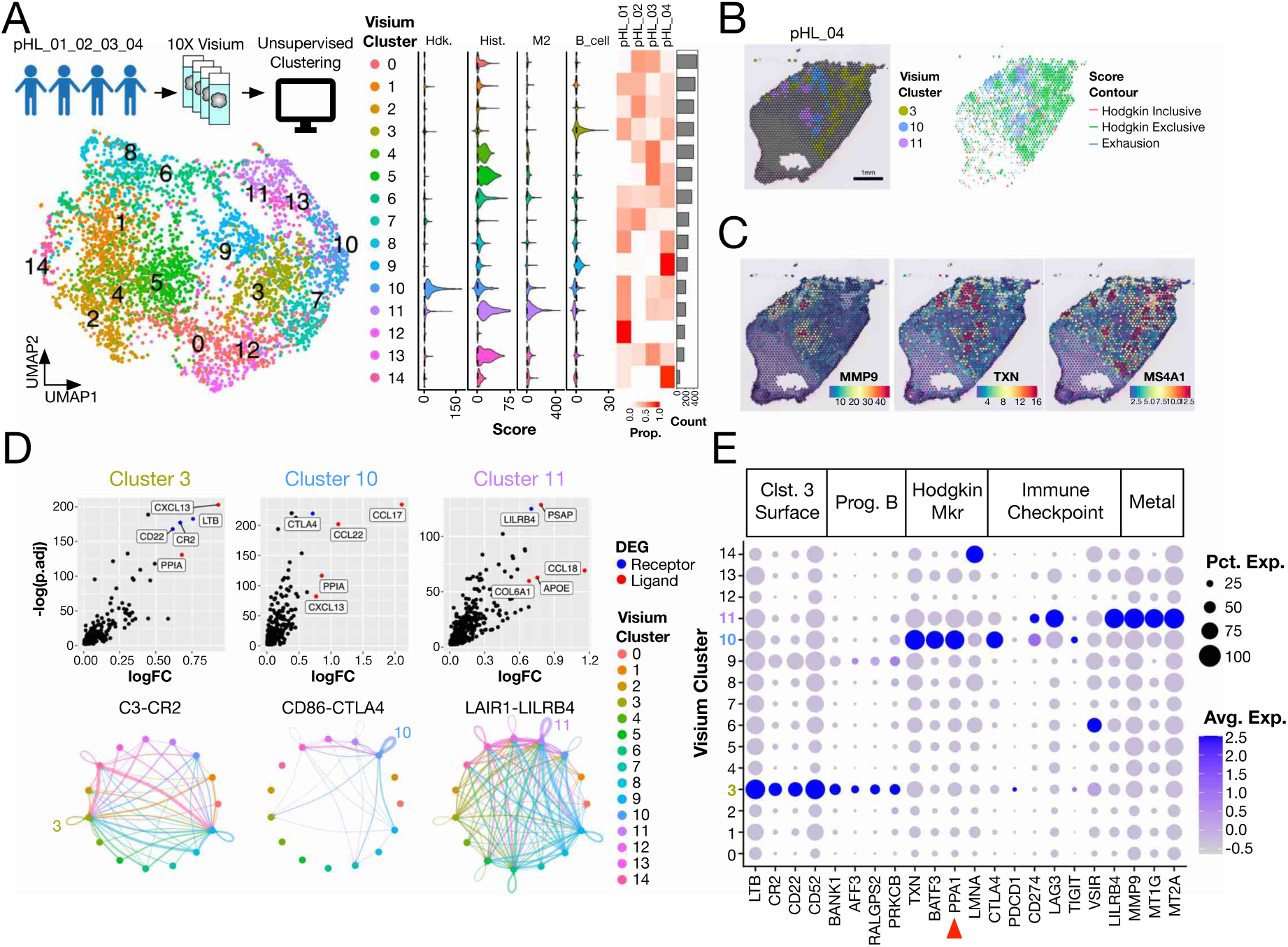
Spatial transcriptomics reveals pHL tumor microenvironment comparments and checkpoint blockade targets: **A.** Characterisation of Hodgkin samples by spatial transcriptomics. (Left) Integration, dimension reduction and unsupervised clustering of Visium spatial dots. (Middle) module scores distribution across clusters. (violin plot from left: Hodgkin, Histiocyte, M2 macrophage-like, B cell). (Right) cell distribution across each sample for different clusters visualised as heat map **B.** Representative Visium slide highlighting locations of clusters of interest (left, cluster3, 10 and 11) and contour plot of module scores calculated by pathologist curated gene sets. Hodgkin inclusive (red contour): *TNFRSF8* (CD30), *FUT4* (CD15), *IRF4*; Hodgkin exclusive (green contour): *MS4A1* (CD20), *CD79A*, *CD19*, *SLC22A2*, *POU2AF1*; Exhaustion (blue contour): *CTLA4*, *LAG3*, *TIGIT*. **C.** Selected spatial gene expression of Hodgkin sample. **D.** Differently expressed gene (DEG) focused on extracellular signalling. (Top row) ligand (in red) or receptor (in blue) transcript upregulated in cluster of interest (from left: cluster 3, 10 & 11). (Bottom row) CellChat visualisation of selected top ligand-receptor pair from above. **E.** Selected gene expression of Visium clusters categorized into surface expression (cluster 3 only), progenitor B, Hodgkin markers, selected T cell immune check point expression and metalloproteins.

To unveil the underlying cause of such TME arrangement, we identified differentially expressed genes (DEGs) for each cluster focusing on extracellular signaling ligand-receptor pairs. We found *CR2* (CD21) among the top upregulated receptors by cluster 3 (B cell) with predicted autocrine-like signaling event within the cluster, suggesting active usage of *C3-CR2* signaling. Similar approach revealed activity of *CTLA4* and *LILRB4*, a previously reported immune checkpoint on myeloid cells (*73*), signaling present in cluster 10 (Hodgkin) and 11 (M2-Mq), respectively (**Figure 6D**). These findings suggest that more than one immune checkpoint is at play in the pHL TME and may be targeted separately. Indeed, we found expression of completely different sets of immune checkpoints by cluster 10 (Hodgkin) and 11 (M2-Mq) with the former specifically expressing *CTLA4* and the latter with high expression of *LAG3* and *LILRB4* (**Figure 6E**). Immunofluorescence microscopy showed *LAG3* expressing cells with or without co-expressing *CD3* in Hodgkin TME, while *PD1* exclusively found in *CD3* positive cells but not *LMNA*, validating the marker gene distribution (**Figure S8C-E**). We detected several genes related to metal metabolism (*MMP9, MT1G, MT2A*) highly expressed by cluster 11 (M2-Mq), providing insights of its interaction with extracellular matrix. Among top marker genes of cluster 10 (Hodgkin), we found *PPA1* (**Figure 6E**), a gene encoding inorganic pyrophosphatase that recently emerged as a novel target for several types of cancer (*74–76*), alongside canonical Hodgkin markers like *TXN* and *BATF3*. As for cluster 3 (B cell), *LTB* was upregulated, indicating high activity in *de novo* genesis of lymphoid structure in this region. Interestingly, we also found top progenitor-like B cell marker genes specifically expressed by cluster 3 (B cell) but not by other clusters, suggesting the presence of progenitor-like B cells outside Hodgkin niche (**Figure 6E**). In summary, spatial transcriptomics revealed a characteristic sandwich-like pHL TME consisting of progenitor-like B cell, exhausted T cell infiltrated Hodgkin niche, and M2-Mq layers, each with distinct biological activity that could be targeted in different ways opening up for next-generation combination approaches for immune checkpoint therapy (*77*).

### Relapsed Hodgkin cells are myeloid-like and maintain PPA1 expression

Current understanding of HL development starts with transformation of pre-apoptotic GC B cells into proliferative, mononuclear Hodgkin cells, which in turn give rise to multinuclear, TME-shaping Hodgkin Reed Sternberg (HRS) cells (*78*). As droplet-based scRNA-seq is not suitable to capture multinuclear HRS cells (*79*), we reasoned that scRNA-seq preferentially captures Hodgkin cells and provides a valuable opportunity to characterize early events in Hodgkin development. Thus, we integrated 4 patient-match pHL samples from Figure 6 (pHL_01∼04, all stage II disease), with a terminal-staged sample acquired upon relapse (pHL_05, stage IV disease) with all pReLy samples, revealing a cluster unique to the pHL sample (between two main B cell clusters, **Figure 7A**), partially overlapping with most stages of B cells and, surprisingly, myeloid cells, indicating certain degree of lineage promiscuity. scVDJ-seq revealed lack of immune receptor expression (**Figure 7B**), ruling out the possibility of normal B nor T cells. We next subclustered pHL samples with inferCNV and combined immune receptor expression, Hodgkin score and CNV score to annotate each subcluster as either Support or Hodgkin (**Figure 7C-D, S9A-E**). As gene expression of HRS cells can resemble histiocytes around them (*66*), the Hodgkin score for the lymphoma subcluster annotation was designed to exclude overlapped transcriptome from macrophage/histiocyte: firstly, we calculated module score of genes documented to be expressed by HRS cells, namely *TXN* and *CCL17* (*63, 80, 81*), followed by subtraction with another module score based on gene known to be exclusively expressed by macrophages: *C1QA*, *LYZ*, *S100A8* (*65, 82, 83*). Of note, *TNFRSF8 (CD30)* expression, the most used gene to identify HRS cells (*84*), was low across all samples and was not included in the Hodgkin module score. In this way, we identified 3 Hodgkin subclones made up of the pHL unique cluster seen in Figure 7A, among which 2 were very small in cell number (∼0.5%) and harbored CNV classically seen in Hodgkin lymphoma like 2p, 12q gain and 13q del (**Figure 7C,E**), aligning with current understanding of Hodgkin cells (*85*). Unexpectedly, pHL_05 was dominated by subclusters with extensive CNV but in general low Hodgkin score, suggesting this sample exhibited a very different transcriptome. Still, most of the common CNV reported in Hodgkin lymphoma such as 9p, 12q, and 17p gain and 13q del was detected in the cluster containing Hodgkin cells (**Figure 7C,E**) (*85*). Interestingly, progenitor-like program was only enriched by non-malignant CNV subclones (**Figure 7D**), again exhibiting lower proportion of cells expressing immune receptors. This agrees with our spatial transcriptomics findings on progenitor-like B cell infiltration.

**Figure 7.**
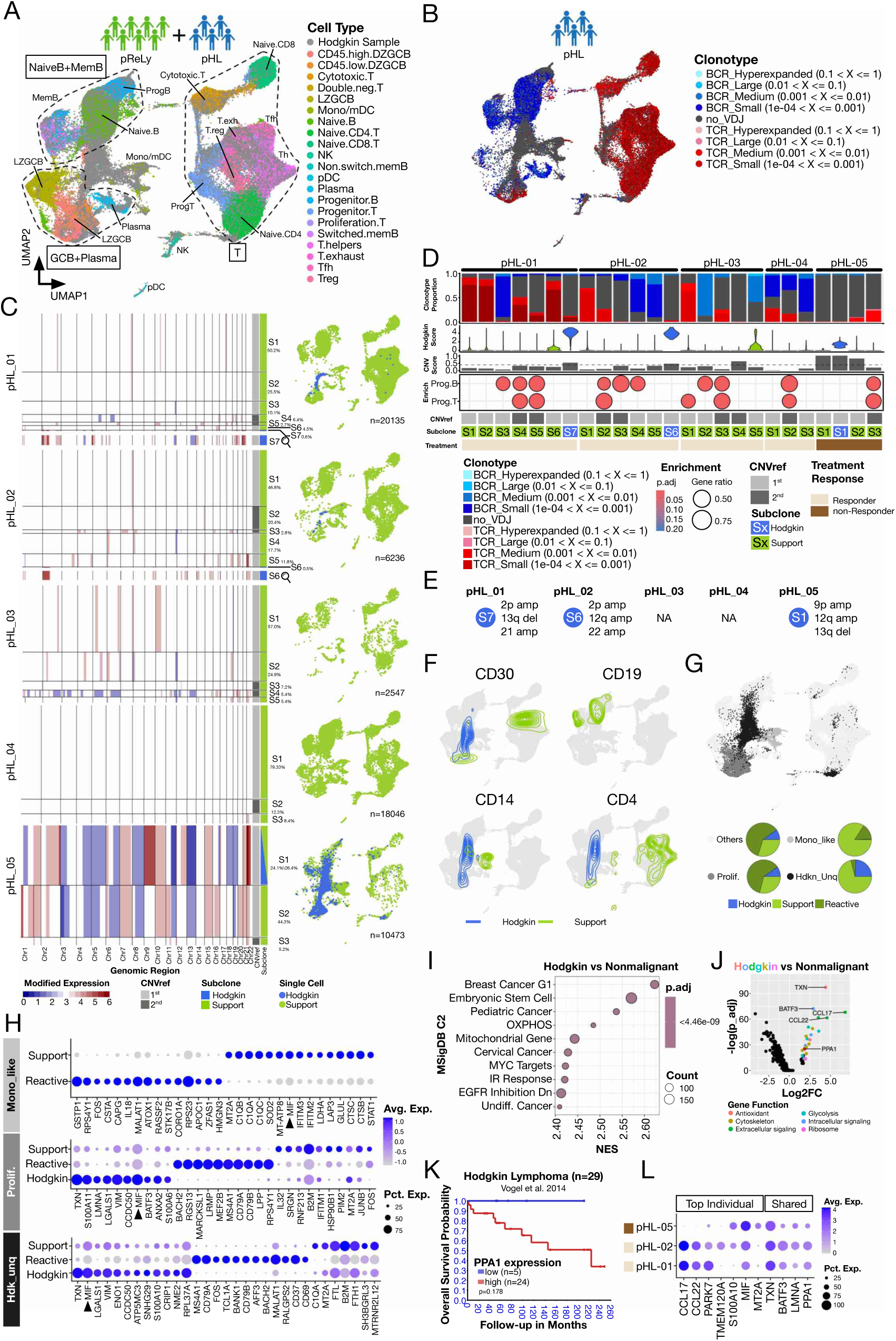
Relapsed Hodgkin cells are myeloid-like and maintain PPA1 expression: **A.** Integration of pReLy (in color) with pHL samples (in grey). **B.** BCR/TCR clonotype of pHL single cells. **C.** (Left) inferCNV subclustering results of pHL using the 1^st^ or 2^nd^ CNVref as reference. (Right) UMAP distribution of CNV subclones eventually annotated as “Support” (in light green) or “Hodgkin” (in royal blue). **D.** Key characteristics of CNV sublcones from **C.** (From top) BCR/TCR clonotype proportion, Hodgkin score, CNV score (dotted line indicates 0.35), over representation analysis using progenitor B/T marker gene sets, CNVref used, and eventual TLBL/Support subclone annotation. **E.** Phylogenetic tree of pHL subclones based on CNV accumulation **F.** Contour plot of Hodgkin or Support single cells with selected gene expression. **G.** Three manually curated cell clusters (Monocyte like, Proliferating & Hodgkin unique) and their composition (pie chart). **H.** Top marker genes of different cell types across different cell clusters defined in G. **I.** Gene set enrichment analysis on ranked transcriptome comparing Hodgkin against non-malignant cells (Reactive + Support) using MSigDB C2 database. **J.** Differently expressed genes (DEG) of Hodgkin comparing to non-malignant cells. Selected genes are highlighted in colors representing different function. **K.** Survival curve of Hodgkin lymphoma patient stratified by *PPA1* expression. **L.** Top individual (left) and shared (right) marker genes of Hodgkin cells from each patient.

Despite that *TNFRSF8 (CD30)* transcript level was low among our samples, its distribution was found to be confined in the pHL unique cluster and exhaustion T cells for Hodgkin and support cells, respectively, suggesting our annotation strategy correctly separated the two compartments (**Figure 7F**). We detected *CD14* and *CD4* that signify monocytes and T helpers in Hodgkin cells in the pHL unique cluster (**Figure 7F**), suggesting lineage promiscuity that has been described in Hodgkin lymphoma (*78, 86*). *CD19* transcript was not detected among Hodgkin cells (top right **Figure 7F**). To characterize the HL TME components, we manually curated three clusters in proximity to the pHL unique cluster: Hodgkin_unique (Hdkn_Unq), monocyte-like (Mono_like) and proliferating (Prolif.). Hdkn_Unq cluster consisted of almost exclusively single cells from pHL sample (**Figure 7G**), underlining the unique transcriptome landscape in pHL TME and the vague transcriptomic boundary between Hodgkin and support cells. Zooming into each curated cluster, we observed: in Mono_like cluster, cells from pHL exhibited distinct expression including complement activities mediated by *C1QA/B/C*, oxidative stress related protein *MT2A*, *SOD2* & *LDHA*, and interferon-stimulated transcription *IFITM2/*3 (**Figure 7H**), all linked to tumor-associated macrophage (TAM) status (*87–91*); as for Hodgkin cells in the other two regions, *TXN* was found as the top marker gene consistently (**Figure 7H**). Interestingly, proinflammatory cytokine *MIF* (*78*) has been found in all three curated clusters by Hodgkin cells or support TAMs (black arrowheads, **Figure 7H**), indicating *MIF* takes the central role shaping pHL TME.

Since the pReLy-defined progenitor-like program was not utilized by lymphoma subclones in pHL, it is likely Hodgkin cells adopt a different precursor program to aid lymphomagenesis. To test this hypothesis, we performed GSEA on ranked genes comparing Hodgkin and non-malignant compartment in a cell-number balanced fashion. Among top enrichment were epithelial cancer-related (like breast and cervical cancers), metabolic and embryonic stem cell terms, suggesting Hodgkin lymphoma utilized a general stemness program common to epithelial originated malignancies (**Figure 7I**). We examined top DEGs from the ranked genes and found *PPA1*, a metabolic enzyme that catalyzes pyrophosphatase hydroxylation, alongside other well characterized genes in HL (**Figure 7J**), and *PPA1* was clinically associated with poor prognosis (**Figure 7K**). Furthermore, *PPA1* expression was detected in Hodgkin cells from all lymphoma subclones (**Figure 7L**), meaning *PPA1* vulnerability was kept upon R/R. Besides, pHL_05 Hodgkin cells had higher expression of myeloid-like genes like *S100A10, MIF* and *MT2A* compared to the non-R/R samples (**Figure 7L**). Taken together, we identified Hodgkin single cells in pHL that depend on *PPA1* expression and were myeloid-like in relapse sample.

### R/R pHL was accompanied by CD74^high^CCL5^+^ CD8 T cells

Since pHL_05 received rituximab previously (**Table 1**), no B cell compartment was found in this particular sample (**Figure 7D**). We therefore focused on the T cell compartment in search of a relapse-specific cell type. We integrated all support T cells from pHL defined by TCR expression in scVDJ-seq and found 12 transcriptomically distinct clusters, among which cluster 7 was uniquely abundant in pHL_05 (**Figure S10A**). Gene expression profile indicated cluster 7 as CD8 T cells expressing high cytotoxic factors like *NKG7* and *GZMA* and exhausting markers *LAG3* and *TIGIT* (**Figure S10A**). As for receptors, we found cluster 7 was void of canonical T cell homing receptors CCR7 and IL7R but exceptionally high in CD74 expression, suggesting that the *MIF-CD74* axis was responsible of recruitment of this cluster (**Figure S10A**). Clonotype analysis revealed cluster 7 harbored several large TCR clones all belonging to pHL_05 (**Figure S10B,C**), indicating weak selection force during clonal expansion. To further characterize these expanding cells, we compared T cells with large TCR clone against other T cells within pHL_05 and confirmed cluster 7 marker genes like *CD8A/B*, *CCL5* and *GZMA* were higher in these expanding T cells (**Figure S10D**), suggesting they were the main effectors of the cluster 7 phenotype. Finally, we examined key cluster 7 gene expression by patient and validated that the phenotype was indeed unique to R/R: we found higher cytotoxic, pro-inflammatory and exhaustion gene expressions in pHL_05, which was rarely found in other samples (NK-like, Granzymes, Cytokine and Exhaustion **Figure S10E**), suggesting a unique T cell cluster in the R/R TME. We confirmed low *CCR7* and high *CD74* expression in T cells through sample-wide comparison (Receptor, **Figure S10E**), pointing to a major shift in T cell recruitment mechanism. In short, we identified a pro-inflammatory T cell subset unique to the R/R TME in pHL and recruited by the *MIF-CD74* axis.

## Discussion

The main challenge in pediatric lymphoma treatment is balancing cure rates against the risk of R/R and the burden of treatment-related toxicity, where R/R is often due to resistance to treatment (*92, 93*). In this study, we identified progenitor-like lymphocytes as a unique trait in children, whose program was enriched in pNHL subclones and likely contributed to R/R especially when becoming an initiating clone for malignant transformation. In pHL the progenitor-like program was found in non-cancerous cells in the TME and was unrelated to R/R. However, Hodgkin cells in R/R were myeloid-like and recruits *CD74^high^CCL5^+^* T cells likely via the *MIF-CD74* axis. Finally, we proposed *MSI2* and *PPA1* as novel vulnerability of pNHL and pHL for further exploitation.

The pediatric-specific, progenitor-like lymphocytes identified in this study are likely to participate in early lymphopoiesis. Indeed, some of the enrichment gene programs like *NOTCH* and *WNT* pathways in progenitor-like lymphocytes have been reported in early lymphopoiesis (*94*), among which *NOTCH2* was documented to promote children specific cell type (*95*). Moreover, a recent study showed that non-cancerous B cells from rituximab treated lymphoma survivor were incapable of maturation (*96*), suggesting *de novo* lymphopoiesis is more complicated than just one step leading to another. Therefore, it is likely that the progenitor-like lymphocytes represent a physiological yet transient compartment that is essential to early lymphopoiesis.

Our data revealed that the progenitor-like program was utilized by lymphoma subclones. In pBL, these subclones showed BCR silencing, traits of chemoresistance, and were often found as the initiating clone in R/R samples. This aligns with findings in diffuse large B cell lymphoma (DLBCL), a closely related GC-derived B cell lymphoma, which became more aggressive upon BCR silencing or extinction (*97, 98*). Such phenomenon has not yet been described in BL, likely because bulk sequencing techniques (*99*) is not suitable to detect the BCR negative compartment which consist roughly 10∼30% of a sample (this study) as compared to up to 65% in DLBCL (*97*). In pTLBL progenitor-like subclones, TCR silencing was observed in 4 out of 5 patients, and pHL are known to be immune receptor negative (*100*), suggesting that immune receptor silencing is a common trait of lymphoma progression. These immune receptor silenced cells, especially in pNHL, likely represent a long-overlooked compartment under the assumption lymphoma cells always express immune receptor. It will be important to better define immune receptor negative subclones to identify markers for pathology evaluation and to identify new treatment modalities for R/R subclones (*28, 101, 102*).

In pBL and pTLBL, lymphoma cells that underwent immune receptor silencing had higher expression of *MSI2*, an RNA binding protein that has been reported to maintain hematopoietic/leukemic stem cells and drive chemoresistance (*34, 103, 104*). Emerging evidence suggests that *MSI2* plays an important role in lymphoma as well, and inhibition of *MSI2* reduces growth of T acute lymphoblastic leukemia in a xenograft model (*105, 106*). When searching for genes complementarily expressed to *TUBB*, *MSI2* was among the top, alongside with enrichment of gene ontology terms that we detected in progenitor-like B and T cells; heightened BCR pathway and stem-like terms, respectively. These lines of evidence place *MSI2* in the center of lymphomagenesis and chemoresistance.

Identification of single Hodgkin cells allowed us to compare transcriptome changes of this early compartment upon relapse, including extensive CNV alteration, a switch to a myeloid like phenotype, and high expression of *PPA1*, an emerging target against several types of cancer (*75*), including colorectal (*74*), lung (*107*), and gastric cancer (*108*). Moreover, Hodgkin cells were enriched in genes related to oxidation phosphorylation (OXPHOS) (**Figure 7I**), a metabolic process where *PPA1* plays a key role. Given that Hodgkin cells are considered a precursor of Reed-Sternberg cells, this unique metabolic vulnerability may provide future treatment benefit. Additionally, we found *CD74^high^CCL5^+^* T cell infiltration in relapsed pHL TME that is well known to respond to *MIF*, an inflammatory cytokine in autoimmune disease like inflammatory bowel disease (*109*), providing new insight to disrupt relapsed HL.

In conclusion, we thoroughly characterized several main types of pediatric lymphoma at single-cell level. Our data reveals chemoresistant progenitor-like pNHL subclones undergo immune receptor silencing, express *MSI2* and, when becoming the initiating clone, is associated with R/R. As for pHL, Hodgkin cells adopt myeloid-like status for relapse and recruit *CD74^high^CCL5^+^* T cell while maintaining *PPA1* as vulnerability.

## Materials and Methods

### Study design and Patient recruitment

The “Pediatric lymphoma” study (EPM: 2021-01381) received ethical approval from the Swedish Etikprövningsmyndigheten in 2021. The recruitment opened in 2021 and recruited patients from Karolinska University Hospital. All children and adolescents (below 18 years of age) with pediatric lymphoma were eligible to enroll at the time of diagnosis upon giving informed consent. Patients with clinically suspected lymphoma who underwent a biopsy or surgery as part of the standard of care were eligible for enrollment into the study which recruited patients from May 2021 to December 2025. The study was performed in accordance with the World Medical Association Declaration of Helsinki. The Code of Ethics of the World Medical Association (Declaration of Helsinki) for human samples was followed. The guardians and patients have provided written consent after receiving oral and written study information.

### Sample collection

All samples were obtained as surgical or core needle biopsies from Karolinska University Hospital, Stockholm, Sweden. All lymphoma specimens were sampled at primary diagnosis except for pHL_05 which was samples at first relapse. Histopathological evaluation was performed by specialist hematopathologists to determine the diagnosis for the different lymphoma subtypes according to the current WHO classification system, as well as for reactive lymph nodes. 8 pediatric reactive lymph node (pReLy) specimens of which one was sampled twice (pReLy_01, 01b), 3 pediatric Burkitt lymphoma (pBL) specimens, 5 pediatric T cell lymphoblastic lymphoma (pTLBL) specimens, 5 pediatric classical Hodgkin lymphoma (pHL) specimens, 1 pediatric anaplastic large cell lymphoma (pALCL), 1 pediatric primary mediastinal large-B-cell lymphoma (pPMBCL), and 2 adult reactive lymph nodes (aLN) were anonymously included in this study and patient characteristics are summarized in the Supplementary tables S1 and S2.

### Single-cell preparation

Samples were mechanically dissociated straight after tissue collection and strained to obtain a single cell suspension. The freshly isolated single-cell suspension used for scRNA-Seq was washed twice with 1x phosphate-buffered saline (PBS) and Dead Removal kit (Miltenyi Biotec #130-090-101) was used to eliminate dead cells. The final concentration of the single cell suspension was adjusted to 900 cells/µL in 1x PBS with 0,04% bovine serum albumin (BSA) following 10X Genomics recommendations. Leftover cell suspensions were cryopreserved as single-cell suspensions in complete RPMI with 10% fetal bovine serum (FBS) plus 10% dimethyl sulfoxide (DMSO) and stored in liquid nitrogen for further analysis.

### Single-cell RNA and VDJ library construction and sequencing

We used the Chromium X instrument, Chromium Next GEM Chip K and the Chromium Next GEM Single Cell 5’ Reagents Kits (V2) to prepare individually barcoded single-cell RNA-Seq libraries and VDJ-Seq libraries following the manufacturer’s protocol (10X Genomics). For quality control and to quantify the library concentration, we used the BioAnalyzer High Sensitivity DNA kit (Agilent Technologies) and Qubit High sensitivity kit (ThermoFischer Scientific). Sequencing using dual indexing was conducted on an Illumina NextSeq machine, using the 150-cycle High Output kit. Sample demultiplexing, barcode processing, and single-cell 39 gene counting were performed with the Cell Ranger Single Cell Software Suite CR2.0.1.

### Bioinformatic pipeline for single-cell RNAseq

For each sequenced scRNAseq library, gene count matrix was generated with 10X CellRanger pipeline (6.1.2) with 2020-A GRCh38 (https://cf.10xgenomics.com/supp/cell-exp/refdata-gex-GRCh38-2020-A.tar.gz) and CR 7.1 (https://cf.10xgenomics.com/supp/cell-vdj/refdata-cellranger-vdj-GRCh38-alts-ensembl-7.1.0.tar.gz) as gene expression and VDJ references, respectively. All quality control and analytic steps were carried out using Seurat package 4.0.5 unless indicated otherwise.

Quality control steps were carried out as follows. First, immune receptor variable genes including IGV, IGD, IGJ, TRV, TRD, TRJ as well as all HLA genes were excluded. Mitochondrial gene percentage was acquired with ‘PercentageFeatureSet()’ function. Single cells from reactive lymph node and lymphoma samples with more than 5% and 10% of mitochondrial gene were excluded, respectively. For reactive lymph node, unique (n.feature) and total (n.count) gene counts were used to exclude single cells with either feature outside the range 300 to mean value plus standard deviation (sd) times 3, and 1000 to mean value plus sd times 3, respectively. The same process was applied to lymphoma samples with different ranges. (BL n.feature = 50 ∼ mean +10*sd, n.count = 50 ∼ mean+10*sd; TLBL & HL n.feature = 150 ∼ mean + 10*sd, n.count = 200 + 10*sd).

High quality single cells passing the abovementioned steps were subjected to standard Seurat workflow. For each sample, gene counts were normalized with ‘NormalizeData()’ function specifying “LogNormalize” method. Top 2000 variable genes were identified using ‘FindVariableFeatures()’ function with “vst” method. Data scaling was done with ‘ScaleData()’ function. Principle components (PCs) were acquired using ‘RunPCA()’ function with top variable genes identified in previous steps, after which ‘doubletFinder_v3()’ from DoubletFinder R package (2.0.3) was used to predict doublet specifying the first 10 PCs. Predicted doublets were removed immediately after this step.

Data integration was carried out step-wise with sanity checks in between. First, we generate integration by sample diagnoses (namely reactive lymph node, BL, TLBL, HL, PMBCL, ALCL), sequentially utilizing Seurat functions ‘SelectIntegrationFeatures()’, ‘FindIntegrationAnchors()’ and ‘IntegrateData()’. Taking batch effect and biological relevance into account, we compared canonical correlation analysis (CCA), reciprocal principal component analysis (RPCA) and harmony as reduction method and found CCA most suitable to our research scope. After that, we further integrated different sample types to facilitate investigation with different scopes. After each integration, we carried out data scaling and PCA on the “integrated” assay of the object and used the first 30 PCs as inputs for ‘RunUMAP()’ function and ‘FindNeighbors()’ function for uniform manifold approximation and projection (UMAP) dimension reduction and Seurat cluster identification.

Cell type annotation was done on integrated reactive lymph node object only. Briefly, we trained two cell type classifiers with a support vector machine-based algorithm scPred (1.9.2) with training reference originating from PBMCs (*32*) and paediatric tonsils (immunesinglecell.com). Generally, each Seurat cluster was annotated in majority vote-fashion. However, conflicts between two predictors and expert knowledge were resolved manually based on marker gene expression which also enable additional cell type annotation wherever feasible. Information from scVDJ-seq was attached to integrated object using scRepertoire (1.7.0) for exclusion of wrongly annotated cells (i.e. annotated B cells having TCR). The final cell type annotation was used as reference to train a pediatric lymph node specific cell type classifier to aid investigation on integration with lymphoma samples.

### Infer CNV, lymphoma subclone annotation & phylogenetic analysis

To detect possible lymphoma cells that lacked hyperexpanded BCR or TCR, we estimated copy number variation (CNV) of each single cell from BL and TLBL samples with inferCNV (1.8.1) against the 1^st^ CNVref described in Figure 1G, setting analysis mode to “subclusters”. Individual subclusters were then tested with their resemblance with 2^nd^ CNVref signatures. Specifically, resemblance was decided by chromosome gain of the first third of the chromosome 10 and loss of the whole chromosome 19. To reveal the true CNV under the 2^nd^ CNVref signature, subclusters identified resembled 2^nd^ CNVref then underwent a second round inferCNV using 2^nd^ CNVref as reference.

CNV subclusters were annotated as either lymphoma or support subclones based on the following steps: for pBL, each subcluster is assigned a CNV score defined as standard deviation of “modified expression” across its full transcriptome. Subclusters with CNV score above cut-off (empirically set to 0.31) are annotated as lymphoma subclones. If a lymphoma subclone defined in this way has low *MYC* expression, it is overwritten as support subclone. If a support subclone upon this step had more than 50% of cells possessing hyperexpanded BCR, it is overwritten as lymphoma subclone; if hyperexpanded BCR is less than 50%, cells with hyperexpanded BCR are separated into a new lymphoma subclone. Lastly, cells with TCR expression are discarded from lymphoma subclone. For pTLBL, subclones with CNV score greater than cut-off (set to 0.18) are annotated as lymphoma. Cells with hyperexpanded TCR in support subclones are separated into a new lymphoma subclone. For pHL, subclones with CNV score and Hodgkin score greater than cut-off (set to 0.35 and 0.7 respectively) are annotated as lymphoma subclones.

Phylogenetic trees of lymphoma subclones from each lymphoma sample was inferred based on the assumption of CNV accumulation. As a validation, we detect transcriptome-wide single nucleotide variation (SNV) at single cell level with SComatic (https://github.com/cortes-ciriano-lab/SComatic) and calculated Jaccard distance of SNV sharing between lymphoma subclones. The Jaccard distance was then used to validate major branching events in phylogenetic trees build by inferCNV.

### Phylogenetic analysis on VDJseq

VDJ sequencing data obtained from Cell Ranger pipeline was analyzed using the Immcantation (www.immcantation.org) framework. VDJ genes for each sequence were aligned to the IMGT GENE-DB database using IgBlast v1.22.0 (REF: https://pmc.ncbi.nlm.nih.gov/articles/PMC3692102/). Change-O v1.3.4 (REF: http://pmc.ncbi.nlm.nih.gov/articles/PMC4793929/) was used to identify the predicted germline and group sequences into clonal clusters. Nonproductive BCRs and TCRs were removed and only heavy chain (IGH) and beta chain (TCRB) data were selected for further analyses. Clonotype frequency and CDR3 characterization were performed for all samples (BCRs and TCRs). Clonotype abundances were computed by counting unique clone_id occurrences, and the 20 most frequent clonotypes per sample were selected for downstream visualization. For these dominant clonotypes, we quantified complementarity-determining region 3 (CDR3) lengths from amino-acid junction sequences and generated per-sample CDR3 length distributions. To characterize sequence diversity within the most abundant clonotype (Clonotype 01), amino-acid sequence logos were created with ggseqlogo v0.2 using probability-based residue frequency estimation. In addition, BCR lineage trees were generated for the top clonotype. Hamming distances were calculated among sequences of equal length using *Biostrings v2.76*, and neighbor-joining trees were built using *ape v5.8*, with the tree rooted on the sequence closest to the germline. Mutation burdens relative to germline and clone-level cell counts were mapped onto the tree using *ggtree v3.16.3*, generating sample-specific phylograms.

### Tissue processing for spatial transcriptomics

Pediatric Hodgkin lymphoma samples (pHL_01-04) were embedded in Optimal Cutting Temperature compound (OCT, Sakura Tissue-TEK) on dry ice and stored at −80 °C. OCT blocks were cut with a pre-cooled cryostat at 8µm thickness, and sections were transferred to fit the 6.5 mm^2^ oligo-barcoded capture areas on the Visium 10x Genomics slide. Before performing the complete protocol, sample quality control was performed according to manufacturer’s instructions: RNA extraction to check RNA quality of the samples (Quiagen RNeasy Mini kit #74104; samples accepted with RIN >9) and Visium Spatial Tissue Optimization (10x Genomics) to select optimal permeabilization. The experimental slide with Hodgkin Reed-Stenberg cells and Hodgkin cells was fixed and stained with hematoxylin and eosin (H&E) and the sequence libraries were then processed according to manufacturer’s instructions (10x Genomics, Visium Spatial Transcriptomic).

### Immunohistochemistry

Pediatric Hodgkin lymphoma samples (pHL_01-04) were embedded in OCT and stored at −80 °C for immunofluorescence staining. OCT blocks were cut with a pre-cooled cryostat at 8µm thickness. Sections were blocked with 5% fetal bovine serum for 1 hour. The sections were stained with CD30 (#BAF1028, R&D systems), CD3 (#300415, Biolegend), PD1 (#AB201825, Abcam), LAG3 (#AB270908, Abcam), LMNA (#AB193903, Abcam), DAPI and mounted with ProLong glass antifade Mountant (#P36984, Invitrogen). Slides were acquired by Zeiss LSM800-Airy using 20X objective. Images were processed with ImageJ software.

### Cell culture

Raji, Jurkat and SUP-T1 cell lines were cultured with complete RPMI1640 with 5, 10, 20 μM of MSI2 inhibitor (Ro 08-2750, # HY-108466, MedChemExpress) or DMSO (vehicle) for 48h. Cells were harvested and labelled for an apoptosis assay with 7AAD (#420403, Biolegend) and Annexin V (#A13201, Invitrogen) and analyzed by flow cytometry. Data was obtained by BD LSRFortessa X-20 flow cytometry machine and analyzed with FlowJo software.

### Flow cytometry

Pathology unit at Karolinska University Hospital provided flow cytometry data for the markers shown in Supplemental Figure 2: CD19, CD3, CD4, CD8, κ and λ Ig chains. From these flow cytometry data it is possible to extrapolate κ/λ and CD4/CD8 ratios as shown in the quantifications.

For flow cytometry data of Figure 3L, pReLy_07 and pBL_03 cells were thawed and dead removal kit (#130-090-101, Miltenyi Biotec) was performed. Then, cells were labelled with LIVE/DEAD Fixable Aqua Dead Cell Stain Kit (#L34966, Invitrogen), CD11b (#101224, Biolegend), CD3 (#317318, Biolegend). PAX5 (#649710, Biolegend), MSI2 (#MA5-57490, Invitrogen), MYC (#MA1-980-AF555, Invitrogen), Ki67 (#350521, Biolegend) and γH2AX (#613420, Biolegend) staining were performed after permeabilization with FOXP3 kit (#421403, Biolegend).

Data were obtained by BD LSRFortessa X-20 flow cytometry machine and analyzed with FlowJo software.

### NSG mice and xenograft

Age- and sex-matched mice were bred and maintained under specific-pathogen-free conditions at the animal facility of the Department of Microbiology, Tumor and Cell Biology, Karolinska Insitutet. All mice experiments performed were approved by the Stockholm North Animal Etics Committee (permit #05294-2023).

1×10^6^ Raji cells were mixed with Matrigel (#356231, Corning) in 1:1 ratio and injected subcutaneously into NSG mouse flank. Starting from day 10, 10 mg/Kg mouse of MSI2 inhibitor (Ro 08-2750; #HY-108466, MedChemExpress) or DMSO control were injected intratumorally every two days. Tumor growth was assessed by digital caliper every 2 days until they reached 1 cm^3^. Statistics were performed using GraphPad Prism version 10. Values were considered statistically significant whether the probability (P) values were equal or below 0,05 (*), 0,01 (**), 0,001 (***) or 0,0001 (****).

## Data and code availability

Codes to reproduce this study are stored on Github (https://github.com/Westerberg-Lab). Annotated single cell objects are stored on European Genome-Phenome Archive (https://ega-archive.org/) as Seurat object with pending accession number.

## Supporting information

All suppementary figures

## Acknowledgement

We are grateful to all patients and their families at the Pediatric Oncology Unit at Karolinska University Hospital for supporting our study. We thank Peri Noori, head of the Center for Molecular Medicine single cell platform at Karolinska Institutet. We thank Gabriela Prochazka, Elisa Basmaci, and Johanna Sandgren at the Childhood Tumor Bank, Karolinska University Hospital and Karolinska Institutet. The data handling was enabled by resources provided by the National Academic Infrastructure for Supercomputing in Sweden (NAISS), partially funded by the Swedish Research Council through grant agreement no. 2022-06725.

This work was supported by Karolinska Institutet PhD (KID) fellowships to T.Y., R.D., a Cancer Society postdoctoral fellowship to J.R., the Childhood Cancer fund and Stockholm region ALF funding to F.B. and L.S.W., the Swedish Research Council, Cancer Society, Worldwide Cancer Research, Radiumhemmet Research Funds, and Karolinska Institutet to L.S.W. L.S.W. is a Ragnar Söderberg fellow in Medicine and holds a senior research position awarded by the Childhood Cancer fund.

## Author Contribution

F.B., L.S.W. conceptualized the study, T.Y., R.D., F.B., L.S.W. designed the research, T.Y., R.D., G.M.M., M.R.L.B., M.H., C.O., J.R. performed the experiments, R.D., T.Y., G.M.M., M.R.L.B., M.H., C.O. and L.S.W. analysed the data, A.E., F.B. recruited and maintained clinical records of the patients, A.K. performed pathology diagnostics and evaluation, T.Y. R.D., G.M.M. and L.S.W. wrote the manuscript, and all authors edited the manuscript.

