## Supplementary material for "Single cell analysis reveals molecular traits of pediatric lymphoma resistant subclones": All suppementary figures

### Supplementary Figure 1

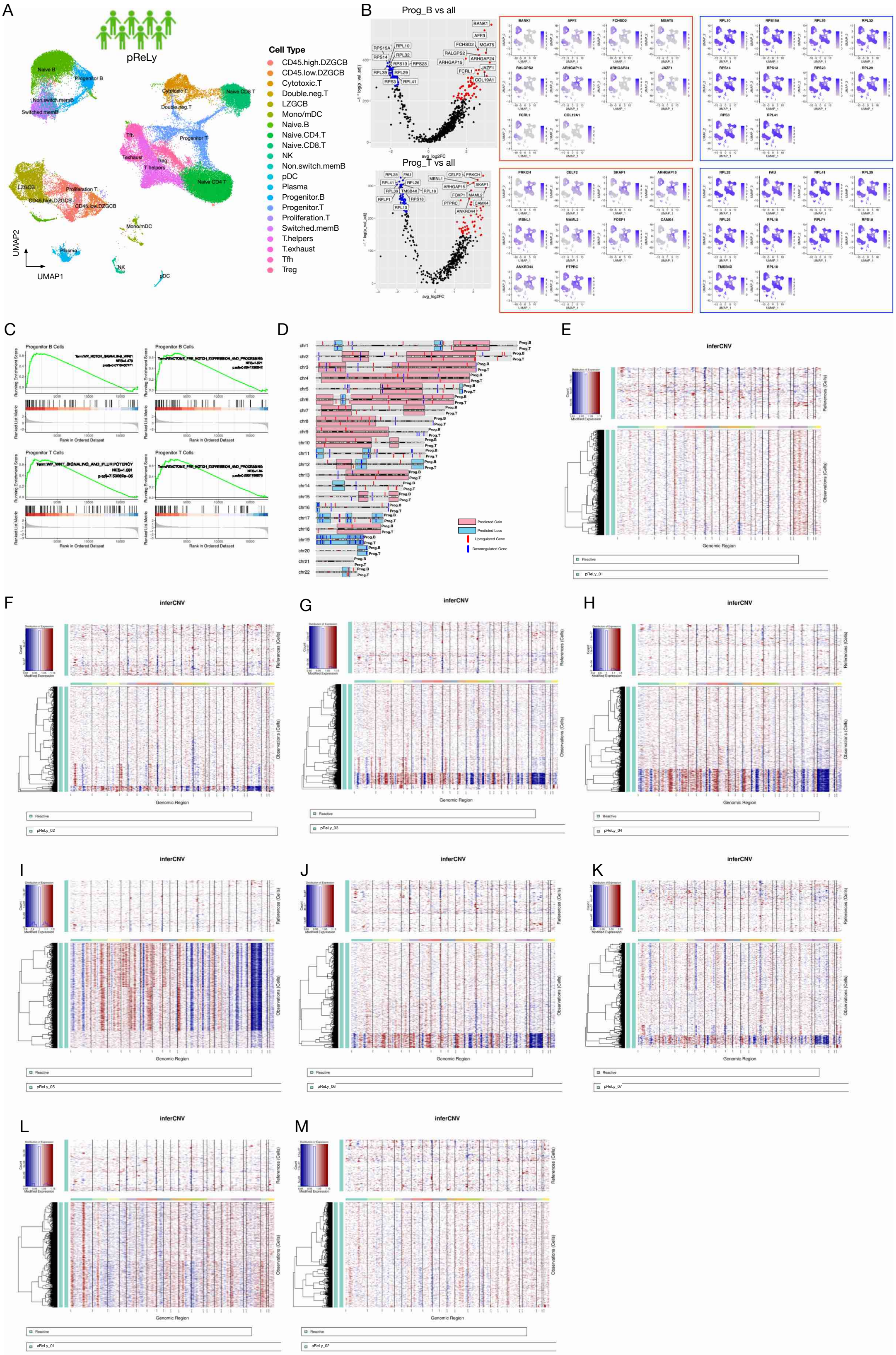

**Supplementary Figure 1. pReLy integrated UMAP and analysis specifications relative to Figure 1.**

UMAP integration of all pReLy specimens for a total of 60,032 cells with color-matched cell types. **B.** (Left) Volcano plot of ProgB/T cells vs. all other cells highlighting downregulation (blue) and upregulation (red) most significant genes. (Right) UMAP representations of those genes. **C.** GSEA of ProgB/T cells focusing on *NOTCH* and *WNT* signalling pathways. **D.** Chromosome position of predicted gain/loss and top up/down regulated genes of progenitor B/T cells**. E-M.** Raw inferCNV output of all a/pReLy in the present study.

### Supplementary Figure 2

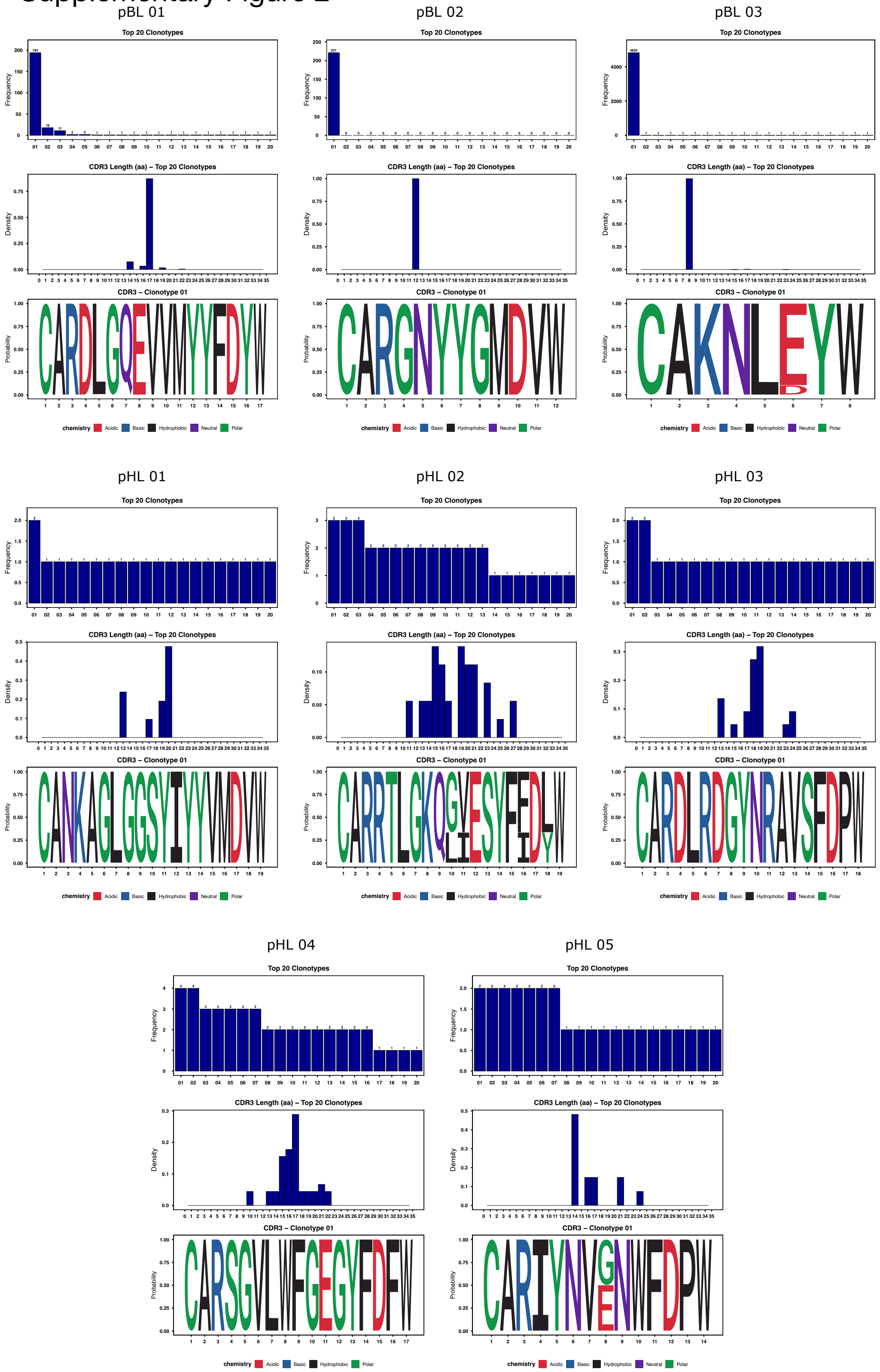

**Supplementary Figure 2. BCR CDR3 analysis of pBL and pHL**

CDR3 protein sequence of largest BCR clone as well as CDR3 length distribution of To 20 BCR clones in each pBL and pHL sample.

### Supplementary Figure 3

A

Gated on CD19+ cells

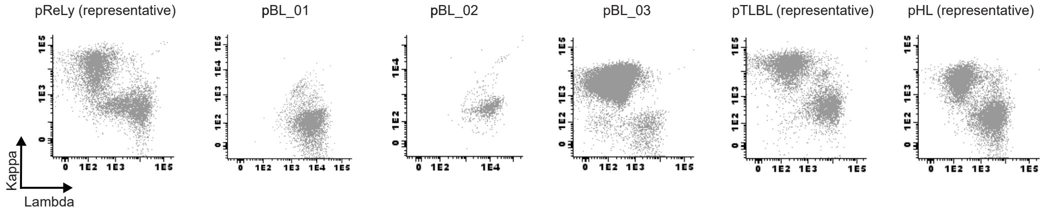


B

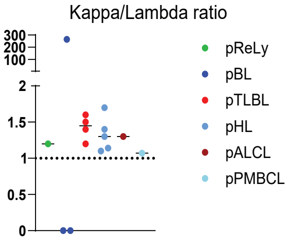


C

Gating strategy for Figure 3L

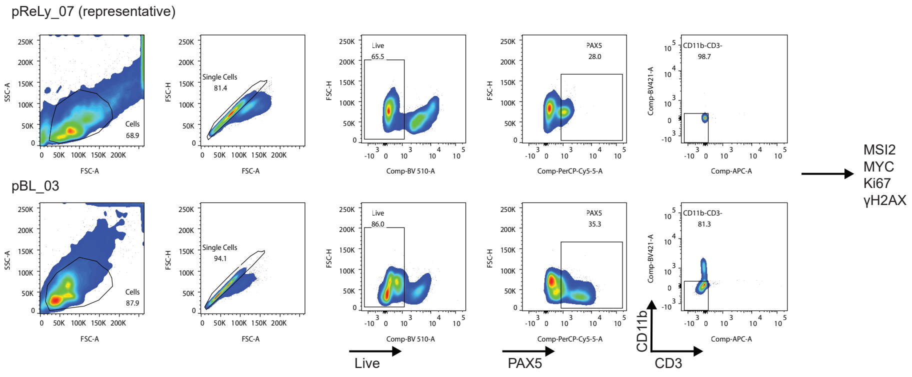

**Supplementary Figure 3. Flow cytometry representations of pediatric tumor specimens.** **A.** Representative flow cytometry data of κ/λ chain usage. One patient for each representative category (pReLy,  pTLL, pHL, pALCL, pPCBCL) and all pBL are shown. **B.** Quantification of κ/λ chain usage referred to panel A. **C.** Gating strategy relative to figure 3L.

### Supplementary Figure 4

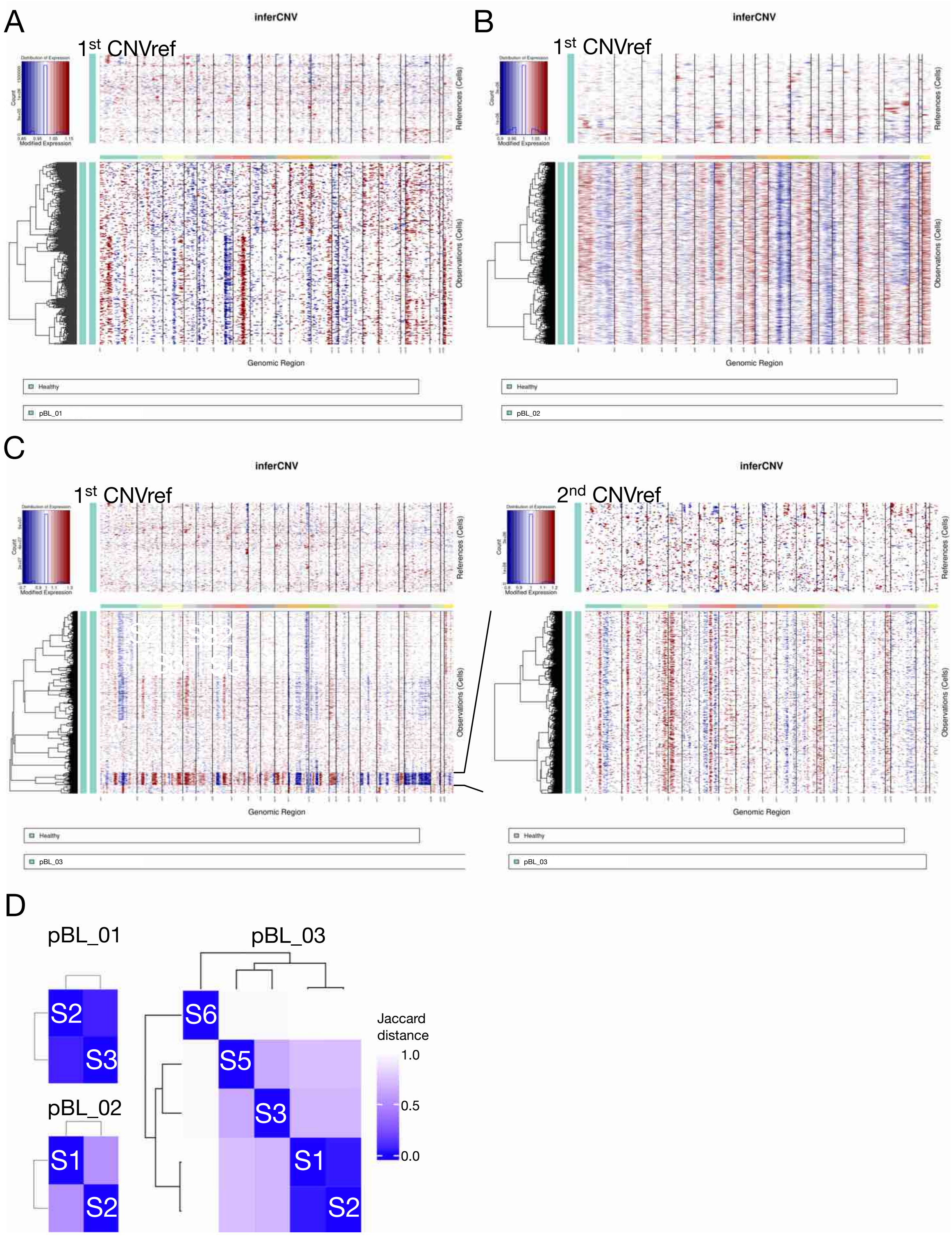

**Supplementary Figure 4. CNV alteration and phylogenetic tree validation of pBL specimens.A.-C.** Raw copy number variation (CNV) profiles inferred from scRNA-seq data are shown as a heatmap. Columns correspond to genomic positions ordered by chromosome, and rows represent individual cells. Red and blue colors indicate relative copy number gains and losses, respectively, compared to reference cells. **C.** CNV pattern of subcluster 5 from pBL_03 resembled $2^{nd}$ CNVref (see method) and was extracted for a second round inferCNV. **D.** Jaccard distance between lymphoma subclones based on shared single nucleotide variation detected by SComatic.

### Supplementary Figure 5

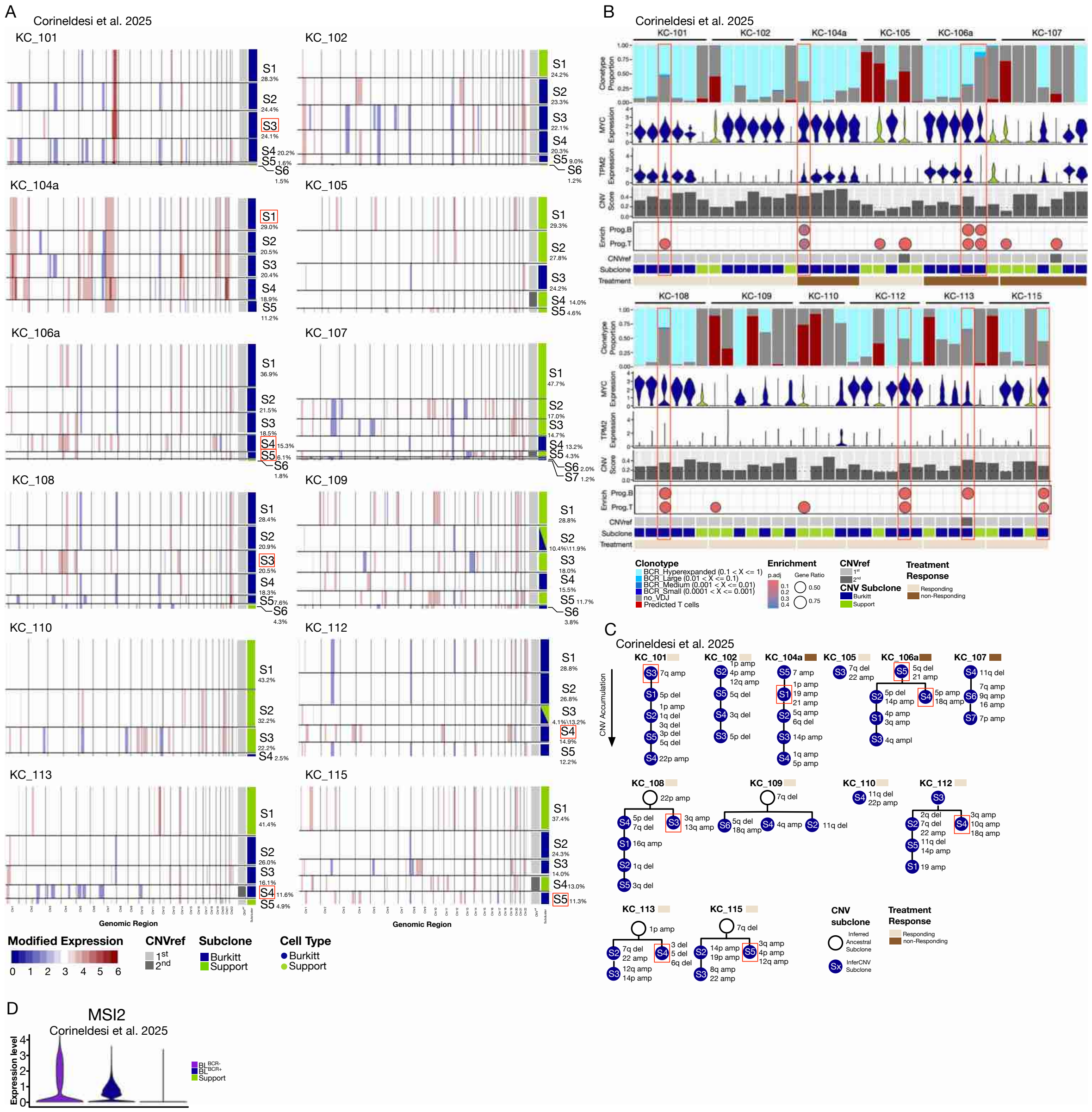

**Supplementary Figure 5. Progenitor-like, BCR silenced pBL subclone found in external data.**

**A.**InferCNV subclustering result of 12 pediatric Burkitt lymphoma sample from Corineldesi et al. 2025. **B.** Key characteristics of each CNV subclones. (From top) proportion of cells with different BCR clone type or predicted T cell status; MYC expression; TPM2 expression; CNV score; enrichment of ProgB/T program; CNVref used; eventual lymphoma/support subclone determination; treatment response. **C.** Phylogenetic tree based on CNV accumulation. **D.** Overall MSI2 expression by lymphoma status and BCR expression.

### Supplementary Figure 6

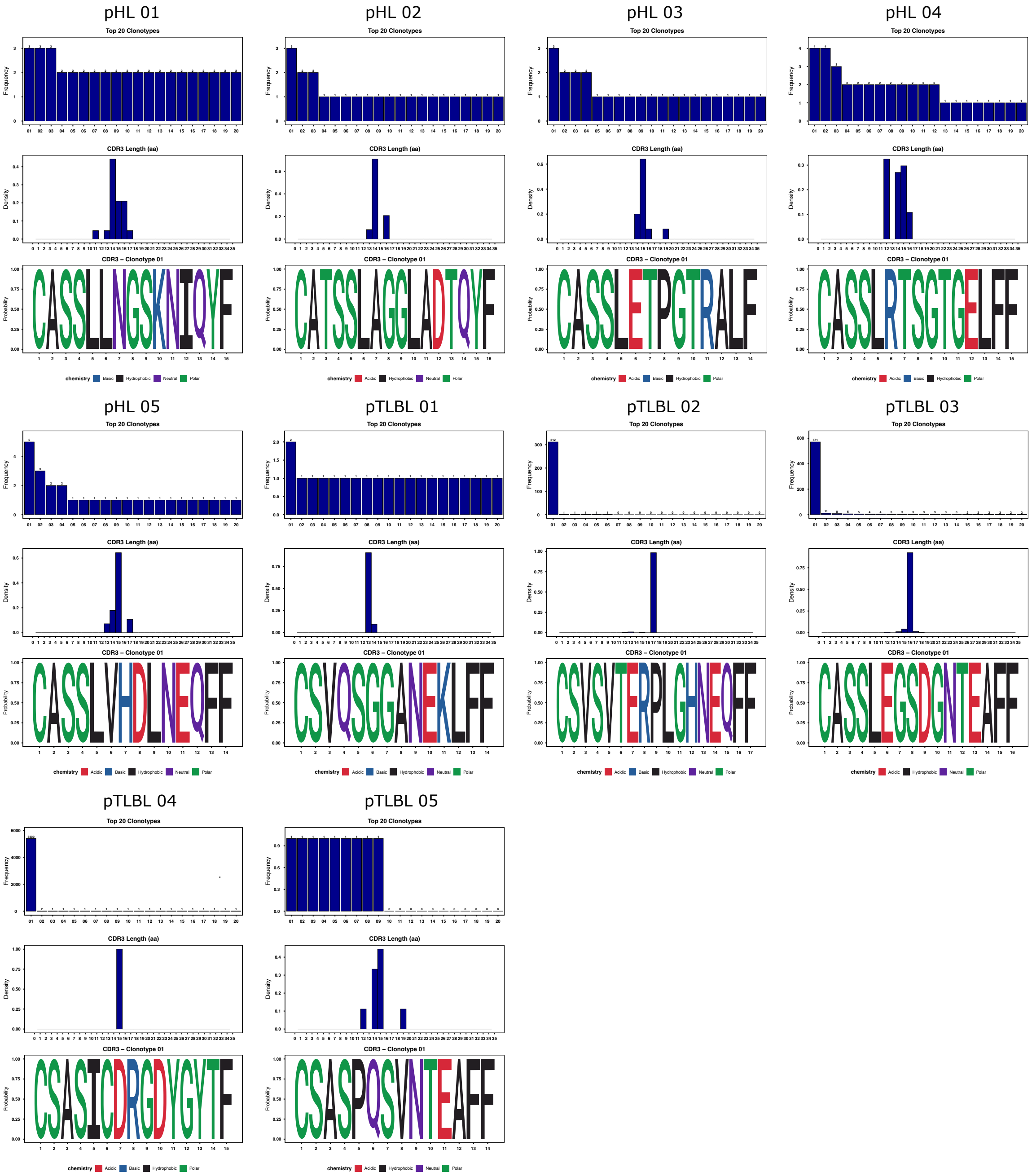

**Supplementary Figure 6. TCR CDR3 analysis of pHL and pTLBL**

CDR3 protein sequence of largest TCR clone as well as CDR3 length distribution of To 20 TCR clones in each pTLBL and pHL sample.

### Supplementary Figure7

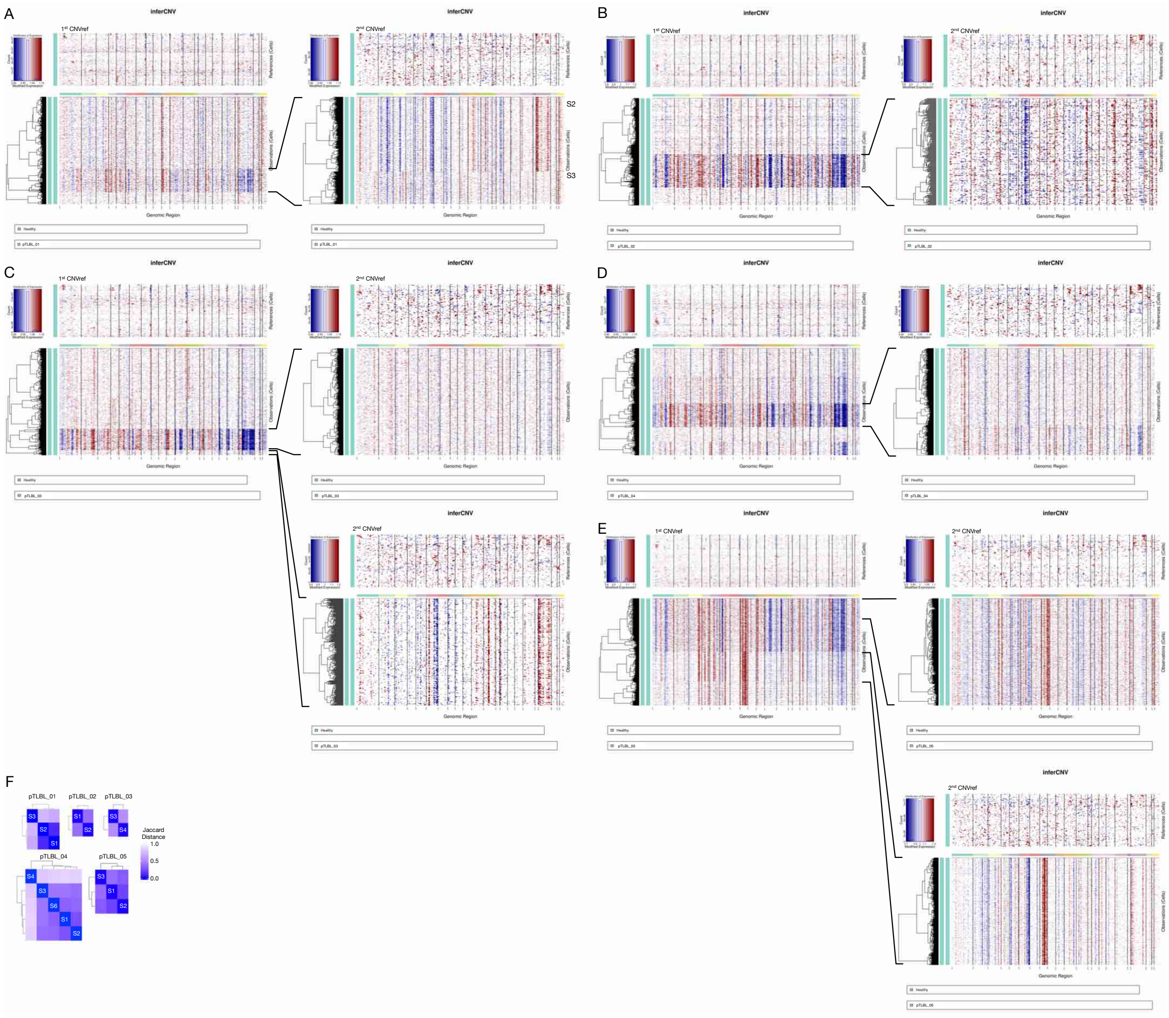

**Supplementary Figure 7. CNV and phylogenetic tree validation of pTLBL patients collected.**

**A.-E.** Raw pTLBL inferCNV result from scRNA-seq data are shown as a heatmap. Columns correspond to genomic positions ordered by chromosome, and rows represent individual cells. Red and blue colors indicate relative copy number gains and losses, respectively, using 1$^{st}$ CNVref (left). Subclusters with CNV pattern resembled 2$^{nd}$ CNVref was extracted for a second round inferCNV with 2$^{nd}$ CNVref (right). **F.** Jaccard distance between lymphoma CNV subclones based on single nucleotide variation (SNV) detected by SComatic.

Supplementary Figure 8

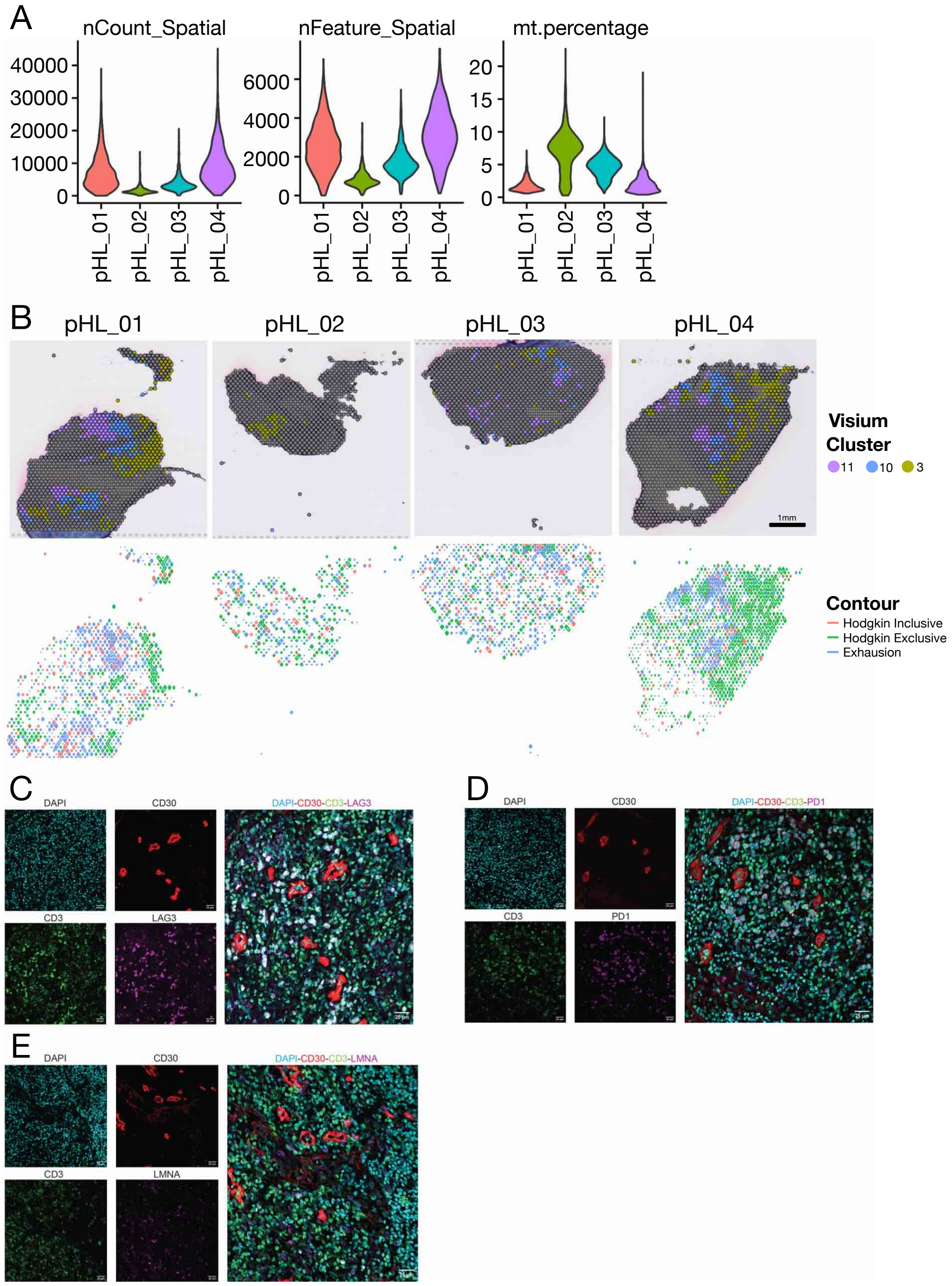

**Supplementary Figure 8. Quality check and validation of spatial transcriptomic findings.** **A.** Spatial transcriptomics sequencing quality information for pHL_01-04 including number of counts, features and percentage of mitochondrial RNA. **B.** (Top) Clusters 3, 10 and 11 highlighted on Visium slides for all pHL samples. (Bottom) Corresponding spatial transcriptomics contour plots highlighting Hodgkin inclusive (red), exclusive (green) and exhaustion (blue) areas. **C.-E.** Representative immunohistochemistry section images of pHL_04. Markers used are: DAPI, nuclei (turquoise); CD3, T cells (green); CD30, HRS cells (red) and exhaustion LAG3/PD1/LMNA (purple).

### Supplementary Figure 9

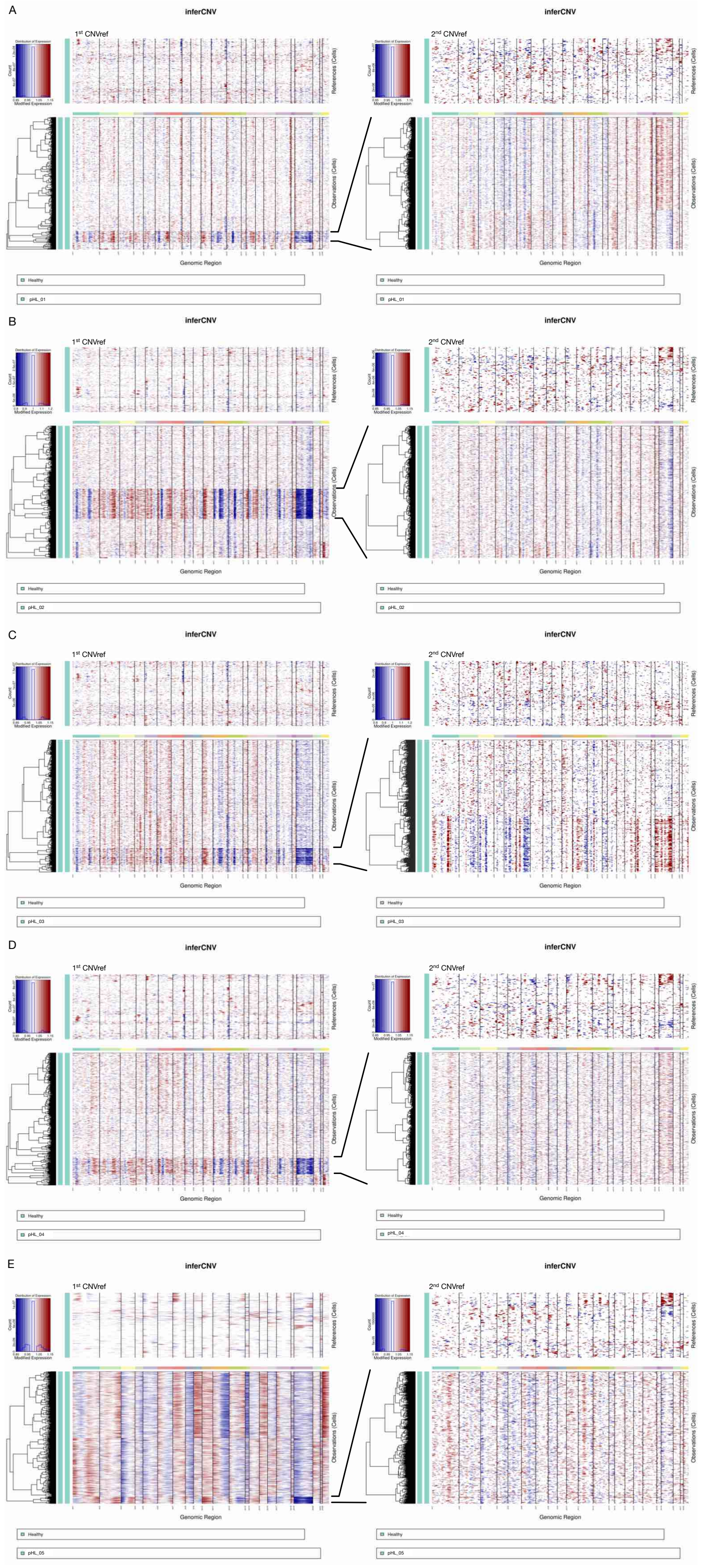

**Supplementary Figure 9. CNV heat map for pHL**

**A.-E.** Raw InferCNV heatmap result of all pHL scRNA-seq. Columns correspond to genomic positions ordered by chromosome, and rows represent individual cells. Red and blue colors indicate relative copy number gains and losses, respectively, compared to the 1$^{st}$ CNVref. 2$^{nd}$ CNVref was used for second round inferCNV on subclusters resembled 2$^{nd}$ CNVref.

Supplementary Figure 10

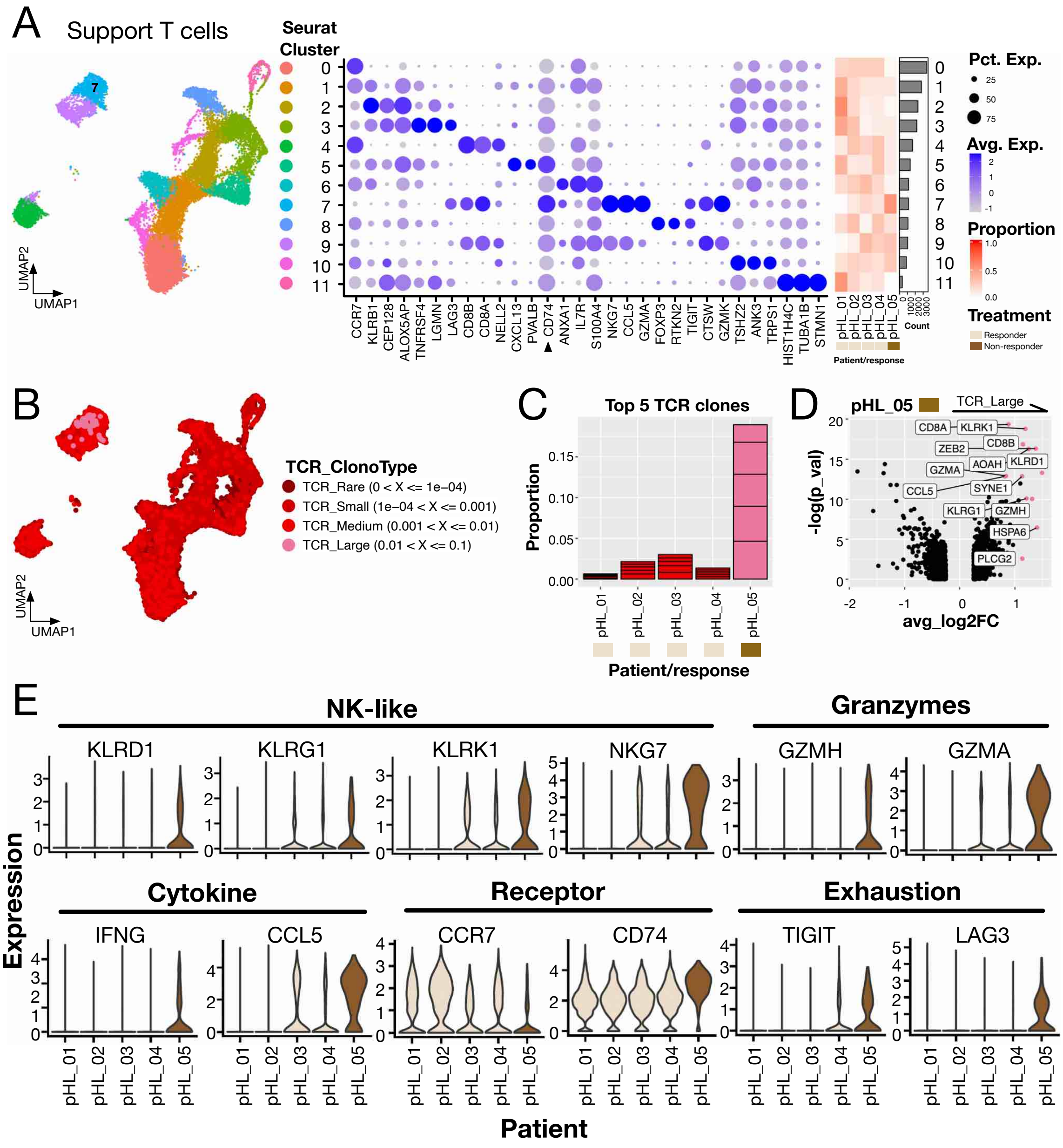

**Supplementary Figure 10. R/R pHL was accompanied by CD74$^{high}$CCL5$^{+}$ CD8 T cells**

**A.** (Left) integration, dimension reduction and bias-free clustering of support T cells from all pHL samples. (Middle) dot plot of top marker genes for each cluster identified in the left. (Right) Heat map showing each cluster's composition by sample. **B.** TCR clone type of each support T cell. **C.** Proportion of top 5 TCR clones from each sample. **D.** Volcano plot highlighting genes upregulated by large TCR clones comparing to other cells from pHL_05. **E.** Violin plot of selected gene expression of biological relevance to pHL_05.
